# Astrocyte molecular rhythm disruption in nucleus accumbens promotes increased binge-like drinking in mice

**DOI:** 10.64898/2026.08.28.747847

**Authors:** Tori Keefauver, Nicole L. Horan, Nicole J. Fairbanks, Lillian Morgan, Anisha Saxena, Ryan W. Logan, Gregg E. Homanics, Sean P. Farris, Marianne L. Seney, Kyle D. Ketchesin

**Author notes:** Email Addresses (in authorship order).

## Abstract

Alcohol misuse is a leading cause of preventable death worldwide. Chronic alcohol is associated with disrupted circadian rhythms, yet molecular mechanisms linking circadian rhythm dysregulation and alcohol consumption are poorly understood. Current FDA-approved treatments for alcohol use disorder (AUD) do not target molecular rhythms or sleep-wake cycles. Mammalian circadian rhythms are regulated by transcription-translation feedback loops that regulate expression of ‘clock genes’ (e.g., *Arntl* encoding for BMAL1). Both human and rodent studies demonstrate associations between clock gene variants and changes in reward-seeking behavior. Evidence suggests astrocytes, non-neuronal brain cells with cell-autonomous rhythms, may regulate both circadian rhythms and reward. In the nucleus accumbens (NAc), a region responsible for modulating alcohol- and reward-related behavior, over 43% of the astrocyte transcriptome is expressed rhythmically. However, no studies to date have investigated roles of NAc astrocyte rhythmicity in regulating alcohol drinking. We used AAV8-Gfap-Cre to functionally ablate molecular rhythms in NAc astrocytes of BMAL1 floxed mice. Two-bottle choice (2BC), drinking-in-the-light (DIL), and drinking-in-the-dark (DID) assessed alcohol drinking. Behavioral assays included locomotor response to novelty, sucrose preference, and social interaction. Disrupting molecular clock function in NAc astrocytes increased binge-like drinking during both DIL and DID paradigms (*d* = 1.44), but not with any other drinking paradigm or behavior. This study carves out a unique role for astrocytes in controlling the temporal organization of reward circuitry and susceptibility to binge-drinking. Future studies will investigate clock-controlled astrocyte mechanisms, such as glutamate uptake and ATP release, that may underlie binge-like drinking behavior.

## 1. Introduction

Alcohol Use Disorder (AUD) is a chronic, relapsing condition affecting 400 million individuals worldwide, and alcohol is a leading cause of preventable death.^1^ The neurobiology of excessive alcohol use is attributed, at least in part, to dysfunctions or disruptions in an individual’s circadian rhythm.^2^ Both clinical and preclinical studies indicate disruptions to sleep-wake cycles often drive people to engage in maladaptive behaviors such as alcohol use.^3^ Some individuals who report chronic alcohol use also report symptoms of disrupted circadian rhythms.^4^ However, prior work has focused on global and region-specific manipulations or genetic polymorphisms underlying the relationship between circadian rhythm dysregulation and alcohol,^3, 5–24^ leaving cell-type specific and molecular mechanisms underexplored. Only three medications are FDA-approved to treat AUD (i.e., disulfiram, naltrexone, and acamprosate), yet none of these treatments were developed to directly address circadian rhythms or circadian symptoms.^25, 26^

Mammalian circadian rhythms are regulated by transcription-translation feedback loops operating on a ∼24-hour cycle. Genes that drive biological rhythms through the transcription-translation feedback loops (e.g., *Clock*, *Per*, *Arntl* encoding for BMAL1) are referred to as ‘clock genes.’^27^ This regulation system for the ‘molecular clock’ is expressed in all mammalian cell types. High expression levels of clock genes form a ‘central clock’ in the suprachiasmatic nucleus, which temporally coordinates all circadian physiology.^28, 29^ Clock gene variants and changes in clock gene expression are associated with significant changes in reward-related behavior in both humans and mice,^15, 30–40^ including alcohol consumption^10, 13, 15–17, 41^ and alcohol-comorbid behaviors.^17, 42^

Evidence across subdisciplines suggests astrocytes may regulate both circadian rhythms and reward processing.^43, 44^ Astrocytes are non-neuronal brain cells that have cell-autonomous molecular rhythms.^45^ Core astrocyte functions including extrasynaptic glutamate uptake and ATP release are regulated by clock genes.^46, 47^ In the nucleus accumbens (NAc), a region responsible for modulating reward-related behavior, we previously determined that over 43% of the astrocyte transcriptome displays rhythmic expression.^48^ We additionally provided evidence supporting a role for NAc astrocyte rhythms in regulating reward.^48^

Prior studies have yet to establish a causal role of astrocyte rhythms in alcohol consumption, circadian timing of alcohol drinking, or alcohol-related behaviors. We hypothesized that alcohol consumption is controlled by astrocyte rhythms in the NAc. To test this hypothesis, we measured alcohol consumption and reward-related behaviors after ablating molecular rhythms in NAc astrocytes.

## 2. Materials and Methods

### 2.1 Animal Care

Adult (6-8 weeks) B6.129S4(Cg)-Bmal1tm1Weit/J (BMFL; RRID:IMSR_JAX:007668) mice (*Mus musculus*) were purchased from the Jackson Laboratory (Bar Harbor, ME). The BMFL strain was originally created on a mixed background [C57BL/6J (B6J) and 129S4/SvJae embryonic stem cells] but is maintained by the Jackson Laboratory on a B6J background using backcrossing. Mice were housed under specific pathogen-free conditions on a 12-hour light/dark cycle (lights on: 07:00, zeitgeber time (ZT) 0; lights off: 19:00, ZT 12) with food [Purina LabDiet PicoLab Isopro RMH 3000 (5P76) (rodent irradiated diet)] and water available *ad libitum*. A total of four homozygous BMFL breeding pairs produced homozygous pups in-house. Homozygous genotypes were confirmed with PCR and gel electrophoresis using primers listed on Jackson Laboratory’s website (**Figure S1**, JAX:007668). Twelve males and 12 females from four litters were used for the virus injected group of alcohol (ethanol), reward, social, and locomotor experiments (Figures 2-3, 5-6). Mice were weaned into cohousing at 21-28 days old, underwent surgery at 6-9 weeks old, and were sacrificed at 33-36 weeks old. Mice were single-housed immediately upon recovery from surgery and remained single-housed until sacrifice. All surgeries were performed during the light cycle (07:00 – 19:00). Adult C57BL/6J (B6J; RRID:IMSR_JAX:000664) mice were purchased from Jackson Laboratory and housed identically to BMFL animals. For the genotype comparison experiment (Figure 4), a cohort of three male B6J, three female B6J, four male BMFL born in-house, and two female BMFL born in-house (from the same breeding pairs mentioned above) were single-housed at nine weeks of age. They remained single housed for three weeks before continuous two-bottle choice (C-2BC) to mimic the length of viral expression and surgical recovery in the virus injected BMFL cohort. All experimental procedures were approved by the Institutional Animal Care and Use Committee of the University of Pittsburgh.

### 2.2 Adeno-Associated Virus (AAV8) Preparation

AAV8-Gfap-Cre-GFP and AAV8-Gfap-eGFP viruses were obtained from the University of North Carolina Vector Core (Chapel Hill NC, RRID:SCR_002448). Specific information about plasmid packaging and promoters used to drive each gene in these vectors is stored by the UNC Vector Core and was previously archived by the National Gene Vector Biorepository. Repository data is not currently available online. Cre and GFP are expressed as a fusion protein. AAV8-Gfap-eGFP was used as a control. NAc expression of Gfap-Cre results in ablation of astrocyte molecular rhythms and loss of astrocyte molecular clock function due to deletion of BMAL1’s functional domain (exon 8; JAX: 007668).^45, 49^ Circadian behavioral rhythms remain intact.^50^ Astrocyte-specific viral knockdown of BMAL1 was previously validated in our colony of BMFL mice after NAc injection.^48^

### 2.3 Stereotaxic Injections and Surgical Procedures

Mice were anesthetized using isoflurane (4% induction, 1.5-2.5% maintenance) and placed in a stereotaxic frame (Kopf Model 942 Small Animal Stereotaxic Instrument with Digital Display Console, Tujunga CA). Heating pads were used during surgery and recovery to prevent hypothermia. Bilateral intracranial microinjections were performed using aseptic surgical techniques. Experimental mice received 1μL per side of either AAV8-Gfap-Cre-GFP (<u>Cre:</u> 5.0×10^12^ vg/mL) or AAV8-Gfap-eGFP (<u>Ctrl:</u> 3.8×10^12^ vg/mL) injected at a rate of 0.2μL/min into the following coordinates relative to bregma: Anterior-Posterior +1.5mm, Medial-Lateral +/− 1.5mm, and Dorsal-Ventral −4.4mm (Kopf Model 1772 Universal Holder, Tujunga CA; Hamilton Glass Syringe, Reno NV). Coordinates were selected using Paxinos and Franklin’s the Mouse Brain in Stereotaxic Coordinates (2001) along with information from previous experiments utilizing the same virus in NAc.^48^ Mice were recovered in a clean cage on a heating pad for 5-20 min until righting reflex and normal ambulatory control returned. Food and water access resumed immediately *ad libitum* and included a liquid diet gel with electrolytes (Clear H2O, Westbrook ME) for the first four days of recovery. Mice received 5mg/kg rimadyl subcutaneously once per day as an analgesic for the first three post-operative days.

### 2.4 Drinking Behavior – Continuous Two-Bottle Choice (C-2BC)

Ethanol and water were administered to mice using 30mL conical tubes (Chemglass Life Sciences Centrifuge Tubes, 30mL, Free Standing, Vineland NJ), rubber stoppers (Fisherbrand One-Hole Rubber Stoppers, size 5.5, Waltham MA), and metal rodent sippers (Braintree Scientific Sipper Tube 1.5” Straight open tube, Braintree MA). Automatic water lixits were removed and replaced with two sipper bottles containing only water four days prior to the start of drinking to acclimate mice to drinking out of sippers. Bottle weights were measured at ZT 0.5 and ZT 12.5 each day during acclimation to minimize neophobia. On the first day of C-2BC, at ZT 0.5, mice were provided with a bottle containing water and a bottle containing 3% ethanol (v/v; Decon Laboratories 200 Proof Ethanol, King of Prussia PA) in water. Every 12 hours (ZT 12.5 and ZT 0.5), bottles weights were measured. Every 48 hours (every other ZT 0.5), bottles were both weighed and replaced with a fresh set. C-2BC was administered according to the timeline in Figure 2, with ramping ethanol concentrations from 3-20%. The bottle containing ethanol was placed on alternating (every 48 hours, corresponding to fresh bottle replacements) left and right sides to control for side preferences. Two empty cages with identical water/ethanol bottle and cage set-ups were used to control for water lost due to cage handling.

### 2.5 Drinking Behavior – Every Other Day Two-Bottle Choice (EOD-2BC)

EOD-2BC was administered according to the experimental timeline in **Figure 2**. Mice received alternating days of two bottles of water and one bottle each of water and 20% ethanol in water. Bottles were weighed and replaced every 24 hours at ZT 8 when diurnal ethanol consumption is at its lowest. All other details regarding cage setup and supplies used are identical to C-2BC.

### 2.6 Drinking Behavior –Drinking in the Light (DIL) and Drinking in the Dark (DID)

DIL and DID were administered according to the experimental timeline in **Figure 2**. Each paradigm lasted for four days. The first three days consisted of two-hour access to one bottle of 20% ethanol (DIL: ZT7-ZT9; DID: ZT14-ZT16). The fourth day consisted of four-hour access to one bottle of 20% ethanol for a binge-like session (DIL: ZT5-ZT9; DID: ZT14-ZT18). Between sessions, mice had access to their typical two bottles of water. All details regarding supplies are identical to C-2BC.

### 2.7 Sucrose Preference Test

Water bottles from home cages were removed and replaced with two 60mL Optimice preference bottles (polypropylene) in a stainless-steel holder (Animal Care Systems Inc., Centennial, CO) filled with water for a 24-hour habituation period. For testing, one bottle was replaced with a 1% sucrose (w/v) solution in sterile drinking water. Starting bottle weights were measured and mice were allowed to drink *ad libitum.* After 24 hours, bottles were reweighed and sides switched for another 24-hour period of *ad libitum* drinking to control for side preference.

### 2.8 Social Interaction Test

Social interaction testing was performed under red light conditions, during the light cycle (ZT 2-5). Prior to testing, mice acclimated to the testing room for 1 hour (ZT 0-1). To measure baseline exploratory behavior and locomotion in the absence of a social target (conspecific), mice were placed in a Plexiglas open-field arena (42 cm × 42 cm × 42 cm, Nationwide Plastics) with an empty small wire animal cage placed at one end near the middle of a wall. Movements were monitored and recorded automatically for 150 seconds (sec) with a tracking system (EthoVision XT 11.0 Noldus Information Technology). At the end of 2.5 minutes, the mouse was removed from the testing arena and the arena cleaned. Next, a same sex conspecific was placed inside the small wire animal cage, and the test mouse was returned to the center of the testing arena. Activity was again measured for 150 sec. For each testing session, time spent in the interaction zone and overall locomotion was recorded. The interaction zone is defined as the half of the open field containing the conspecific. Social interaction ratio was calculated by dividing the time spent in the interaction zone with conspecific by the time spent in the interaction zone when conspecific was absent. Three male and three female conspecifics were rotated throughout testing.

### 2.9 Locomotor Response to Novelty

Light phase testing occurred from ZT 1-4 and dark phase testing occurred from ZT 13-18 under red light conditions. All testing was performed in clear Plexiglas test chambers (Kinder Scientific Smart Cage Rack System; field dimensions: 9.5” × 18.0”) equipped with infrared photobeams measuring horizontal locomotor activity (EthoVision XT, RRID:SCR_000441). Before beginning each session, mice acclimated to the testing room for 1 hour. Sessions lasted 120 minutes (min) and measured total distance traveled (cm) in 5-min bins. Chambers were cleaned between groups with Peroxiguard (active ingredient: hydrogen peroxide). Mice were randomized to testing order, with equal numbers per virus group assigned to each sequence. Twelve mice were tested in the light first and dark second, and the remaining 12 were tested in the reverse order to control for potential order effects.

### 2.10 Euthanasia

Mice not needed for histology were euthanized using carbon dioxide and secondary cervical dislocation. BMFL mice from the surgical cohort were anesthetized with isoflurane (3% for 10-20 minutes) and transcardially perfused with phosphate buffered saline (PBS). Brains were rapidly extracted and snap frozen on dry ice.

### 2.11 Tissue Processing and Sectioning

Samples were frozen at −80°C until sectioning. Brains were embedded in Optimal Cutting Temperature compound immediately prior to sectioning. Fifteen-micron sections containing NAc were obtained using a cryostat (Leica CM1950, Wetzlar, Germany, Leica: CM1950 Cryostat, RRID:SCR_018061) and mounted directly onto coated slides (Superfrost Plus, Fisher Scientific, Pittsburgh PA). Slides were stored at −80°C.

### 2.12 Imaging

Cells expressing the virus were visualized using the endogenous GFP fluorescence; an anti-GFP antibody was not necessary. Frozen sections were fixed in 10% neutral buffered formalin overnight at 4°C, followed by a wash in 1x PBS. Slides were cover slipped using DAPI (NucBlue, Invitrogen, Euguene, OR) and aqueous mounting medium (ProLong Diamond Antifade Mountant, Invitrogen, Euguene, OR). Fluorescent images of viral placements (GFP) were collected at 20x using an Olympus Slideview VS200 slide scanning microscope with OlyVIA software (RRID:SCR_024783). Images were quantified using Qupath: (https://qupath.github.io, RRID:SCR_018257).

### 2.13 Exclusion Criteria

Fluorescent images of viral placements revealed five females with misplaced unilateral injections (left side). These females were excluded from all figures and analyses.

### 2.14 Statistical Analysis

Raw weights of each water bottle (g), solution densities (g/mL), and mouse body weights (g) were used to calculate ethanol consumption (g/kg), total fluid intake (g/kg), and ethanol preference [ethanol(g)/total fluid(g)]. Average grams of water/ethanol lost from control cages each day were subtracted from raw values for each mouse to produce leakage-corrected ethanol and water consumption values. Data was analyzed for effects of each viral condition, sex differences, and interaction effects. Sexes were collapsed when there was no main effect of sex, but sex-split data is analyzed and available in **Figures S2 and S3**. Calculations were completed using Microsoft Excel. Figures and statistical tests were generated using GraphPad Prism (https://www.graphpad.com, RRID:SCR_002798). Data were analyzed using Shapiro-Wilk and Kolmogorov-Smirnov tests for normality (sample-size permitting), two-way ANOVAs, and three-way repeated measures ANOVAs. When repeated measures values were missing due to random errors in data collection, a mixed-effects model using maximum likelihood was fit instead of an ANOVA. The Geisser-Greenhouse correction was applied to repeated measures ANOVAs and mixed models. Two-tailed Welch’s t-tests were used in sex-collapsed, non-repeated measures analyses, with pooled standard deviation used to calculate Cohen’s *d* (effect size). Significant statistical results are reported in the Results. Full statistical results are reported in the extended figure captions (See Supplementary Material, Section II).

## 3. Results

To investigate the impact of astrocyte molecular rhythms on alcohol (ethanol) drinking, BMFL mice received NAc microinjections of adeno-associated virus expressing either a fusion protein containing Cre recombinase with a green fluorescent protein (GFP) reporter, or a control virus expressing only enhanced-GFP (eGFP), both under control of the astrocyte-specific Gfap promoter (**Figure 1**). To assess ethanol drinking and associated reward and circadian behaviors, mice were run through a battery of drinking and behavioral assessments (**Figure 2**): continuous two-bottle choice (C-2BC), every-other day two-bottle choice (EOD-2BC), drinking-in-the-light (DIL), drinking-in-the-dark (DID), sucrose preference test (SPT), social interaction test (SIT), and locomotor response to novelty (LRN).

**Figure 1:**
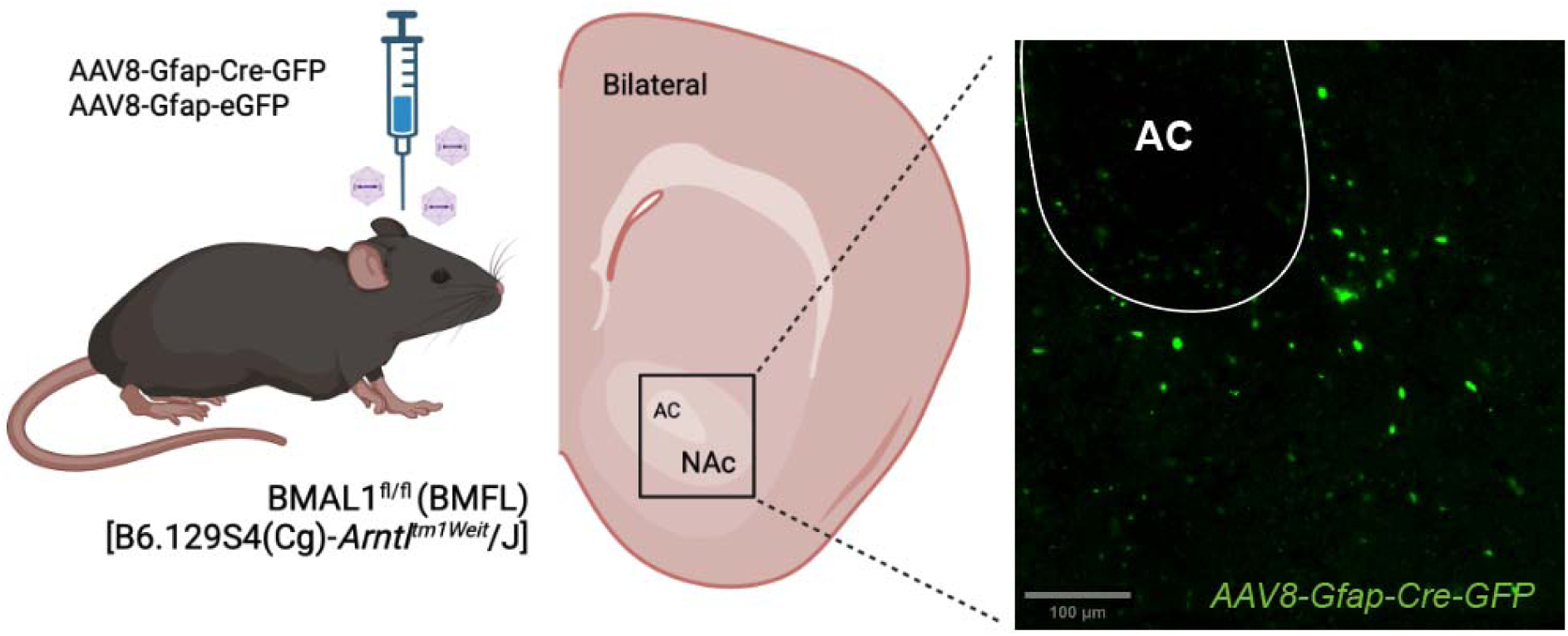
**(A)** Schematic illustrating bilateral stereotaxic injections of either AAV8-Gfap-Cre-GFP or AAV8-Gfap-eGFP virus into NAc of BMFL mice; injections at 1uL/min, coordinates relative to bregma: AP: +1.5, ML: ± 1.5, and DV: −4.4; angle 10°. AC=Anterior Commissure. Created in BioRender. Keefauver, T. (2026) https://BioRender.com/fli9rh2

**Figure 2:**
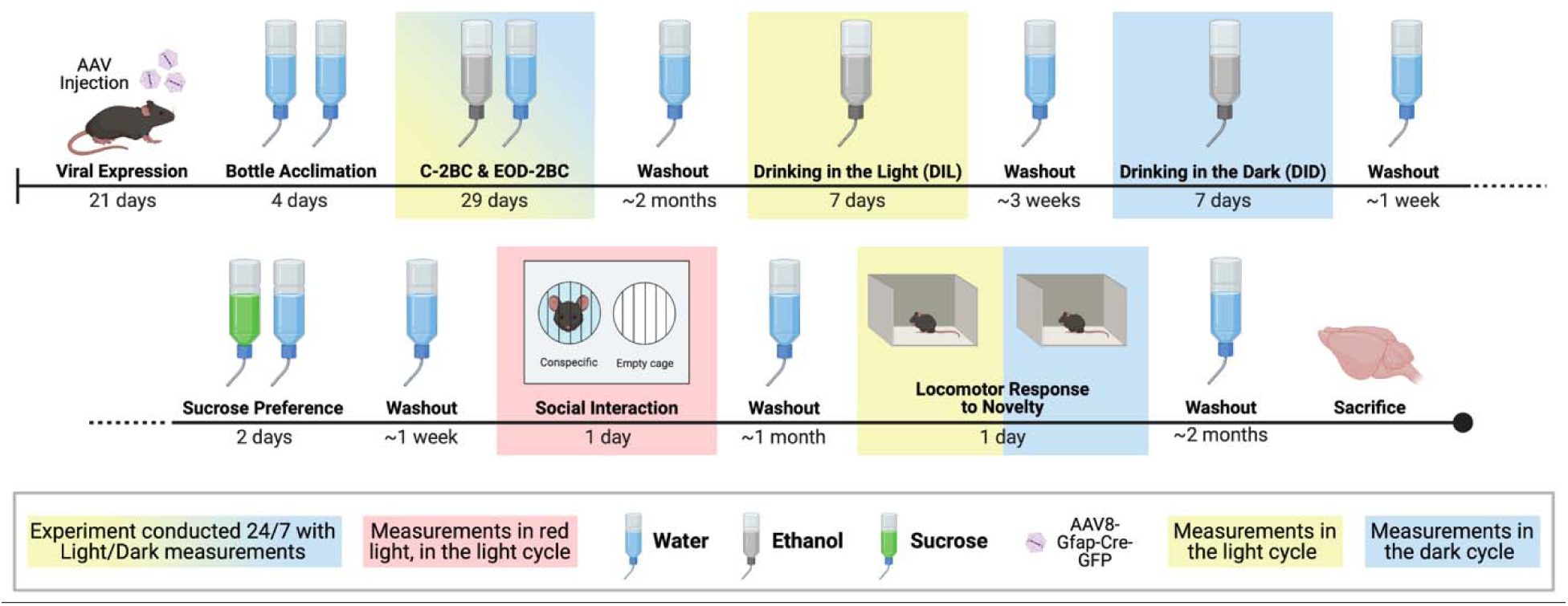
Overview of Experimental Design. Schematic illustrating full experimental timeline for BMFL mice (n=12 males; n=7 females) receiving viral injections. Experiments conducted in the following light conditions: Yellow = light cycle only; Blue = dark cycle only; Yellow/Blue gradient = 24/7 but with separate measurements in light and dark cycles; Red = during the light cycle but under red light conditions. Created in BioRender. Keefauver, T. (2026) https://BioRender.com/r82ixtj

### 3.1 BMAL1 functional ablation in NAc astrocytes does not impact voluntary drinking in continuous or every other day access two-bottle choice

BMFL mice were first tested with a C-2BC assay followed immediately by EOD-2BC, drinking assays validated for measuring home-cage ethanol consumption and preference (**Figure 3A**).^51–54^ Measurements of ethanol consumption for C-2BC were obtained every 12 hours at ZT 0.5 and ZT 12.5 to measure ethanol consumption in the light and dark cycles separately.

**Figure 3:**
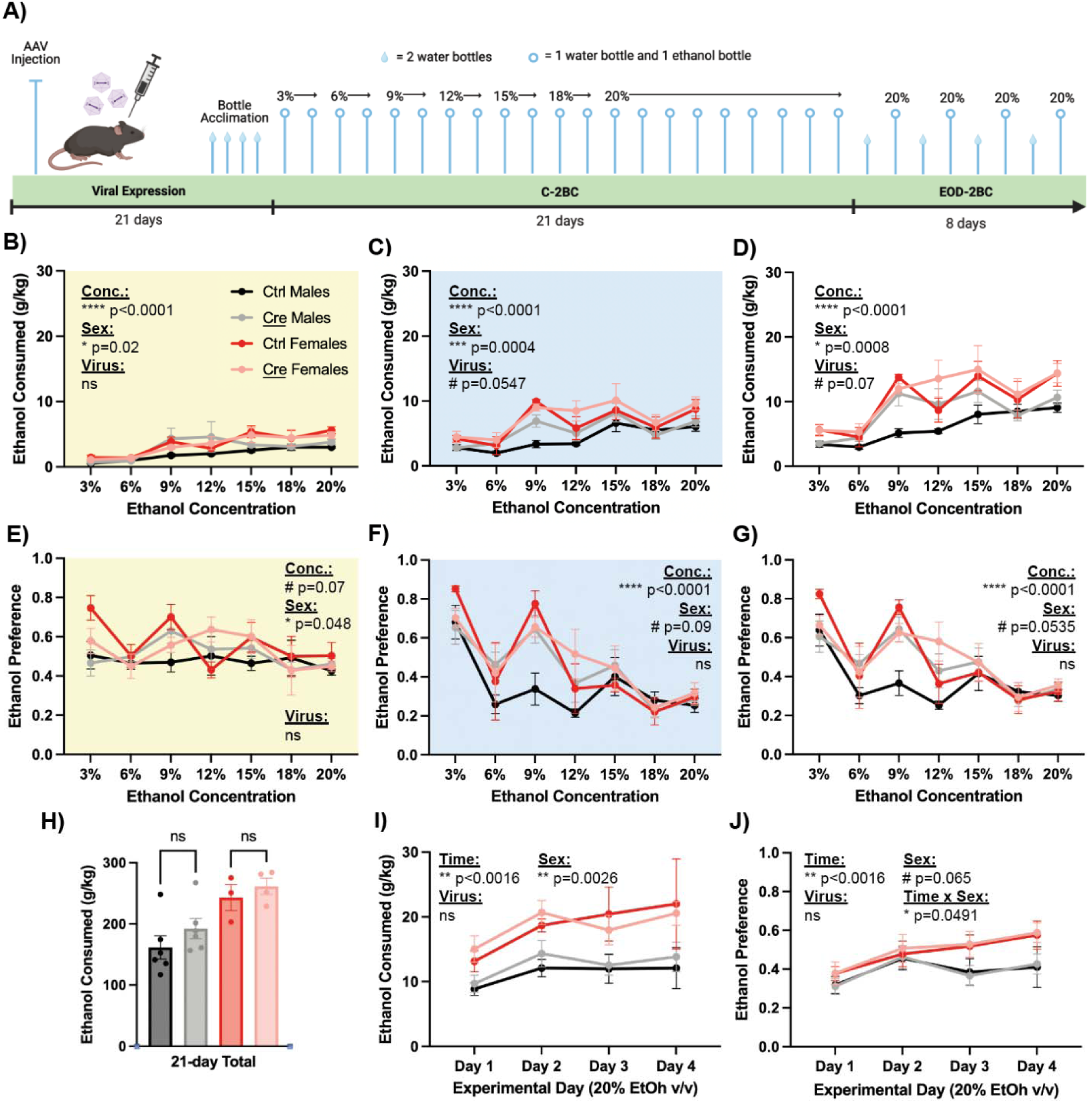
**(A)** Schematic detailing time course for viral injections, bottle acclimation, continuous two-bottle choice (C-2BC), and every other day two-bottle choice (EOD-2BC) experiments in BMFL mice (n=12 males; n=7 females). **(B)** Mice with a loss of BMAL1 function in NAc astrocytes (Gfap-Cre) show no differences in ethanol consumption (g/kg) in the light cycle during C-2BC drinking, and females consumed significantly more alcohol than males. **(C)** During the dark cycle, females still consume more ethanol than males, and there is a trend towards a significant effect of BMAL1 loss on ethanol consumption. **(D)** When data is collapsed into 24-hour periods, results mimic the dark cycle. **(E)** Mice with a loss of BMAL1 function in NAc astrocytes show no differences in ethanol preference in the light cycle during C-2BC drinking, and females preferred ethanol significantly more than males. **(F)** During the dark cycle, there is a significant effect of ethanol concentration on ethanol preference and a trend towards a significant effect of sex. **(G)** When data is collapsed into 24-hour periods, results mimic the dark cycle. **(H)** Mice with loss of BMAL1 function in astrocytes show no within-sex differences in cumulative ethanol consumption across the 21-day task. **(I)** In an EOD-2BC drinking task, mice show main effects only of sex and time on ethanol consumption, with females consuming more ethanol than males, and both sexes increasing their drinking across the 8-days. **(J)** Similar to EOD-2BC ethanol consumption, there is a significant effect of time and a trend towards a significant effect of sex on ethanol preference. There is an additional significant interaction between time and sex, with only females exhibiting a stronger preference for ethanol at the end of the task. Experiments conducted in the following light conditions: Yellow = light cycle only; Blue = dark cycle only. #p<0.1, *p<0.05, **p<0.01, ***p<0.001, ****p<0.0001. Created in BioRender. Keefauver, T. (2026) https://BioRender.com/f4ibihx

In C-2BC, we observed no significant effect of BMAL1 functional ablation in NAc astrocytes on ethanol consumption or ethanol preference; results were consistent when assessed in the light cycle, dark cycle, or collapsed into 24-hour blocks (**Figure 3: B-G**). Cumulative ethanol consumption summed across the 21 days of C-2BC was also not significantly different between virus groups (**Figure 3H**; <u>Virus:</u> F(1,15) = 1.675, p=0.2152). There was a trend towards a main effect of virus on ethanol consumption in both the dark cycle (<u>Virus:</u> F(1, 15) = 4.342, #p=0.0547) and 24-hour blocks (<u>Virus:</u> F(1, 15) = 3.871, #p=0.0679) (**Figure 3: C-D**).

During the dark cycle and 24-hour blocks, there was a main effect of sex on ethanol consumption (g/kg) (**Figure 3: C-D**; <u>Sex_Dark_:</u> F(1, 15) = 20.14, ***p=0.0004; <u>Sex_24hr_:</u> F(1, 15) = 17.69, ***p=0.0008) and a trend towards a main effect of sex on ethanol preference (**Figure 3: F-G**; <u>Sex_Dark_:</u> F(1, 15) = 3.313, #p=0.0887; <u>Sex_24hr_:</u> F(1, 15) = 4.393, #p=0.0535). Females exhibited higher ethanol consumption and ethanol preference compared to males, consistent with prior evidence for basal sex differences in mouse ethanol consumption.^55, 56^ During the light cycle, a significant main effect of sex was observed on both ethanol consumption (**Figure 3B**; <u>Sex:</u> F(1, 15) = 6.605, *p=0.0213) and ethanol preference (**Figure 3E**; <u>Sex:</u> F(1, 15) = 4.637, *p=0.0480), with females greater than males on both measures. There was also a significant main effect of sex on cumulative ethanol consumption (**Figure 3H**; <u>Sex:</u> F(1, 15) = 15.62, **p=0.0013).

In EOD-2BC, we also observed no significant effect of BMAL1 functional ablation in NAc astrocytes on ethanol consumption or ethanol preference (**Figure 3: I-J**). There was a significant main effect of sex on ethanol consumption (**Figure 3I**; <u>Sex:</u> F(1, 15) = 13.03, **p=0.0026) and a trend towards a main effect of sex on ethanol preference (**Figure 3J**; <u>Sex:</u> F(1, 20) = 3.816, #p=0.0649). Females consumed more ethanol and had a higher ethanol preference compared to males, thus remaining consistent with previously established sex differences in basal mouse ethanol consumption.^55, 56^

### 3.2 BMFL mice consume and prefer ethanol the same as sex- and age-matched B6J mice

After observing a relatively low preference for ethanol in BMFL mice (**Figure 3: E-G**), we established a new non-surgical cohort of BMFL mice alongside sex- and age-matched B6J mice to assess the impact of BMFL genotype on ethanol consumption. No viruses were delivered to these mice, and the B6J comparison mice were bred in-house under the same developmental conditions. Mice were single-housed for three weeks to mimic surgical recovery and tested using an identical C-2BC paradigm (**Figure 4A**). There were no differences in ethanol consumption or ethanol preference due to genotype (**Figure 4: B-C**). There was again a basal sex difference in ethanol consumption (Sex: F(1, 8) = 11.81, **p=0.0089), with females consuming significantly more ethanol (**Figure 4: B-C, Figure 3: B-H**). These results suggest no effect of the mixed genetic background of the BMFL on ethanol consumption, and no differences in BMFL vs B6J ethanol consumption in our animal facility.

**Figure 4:**
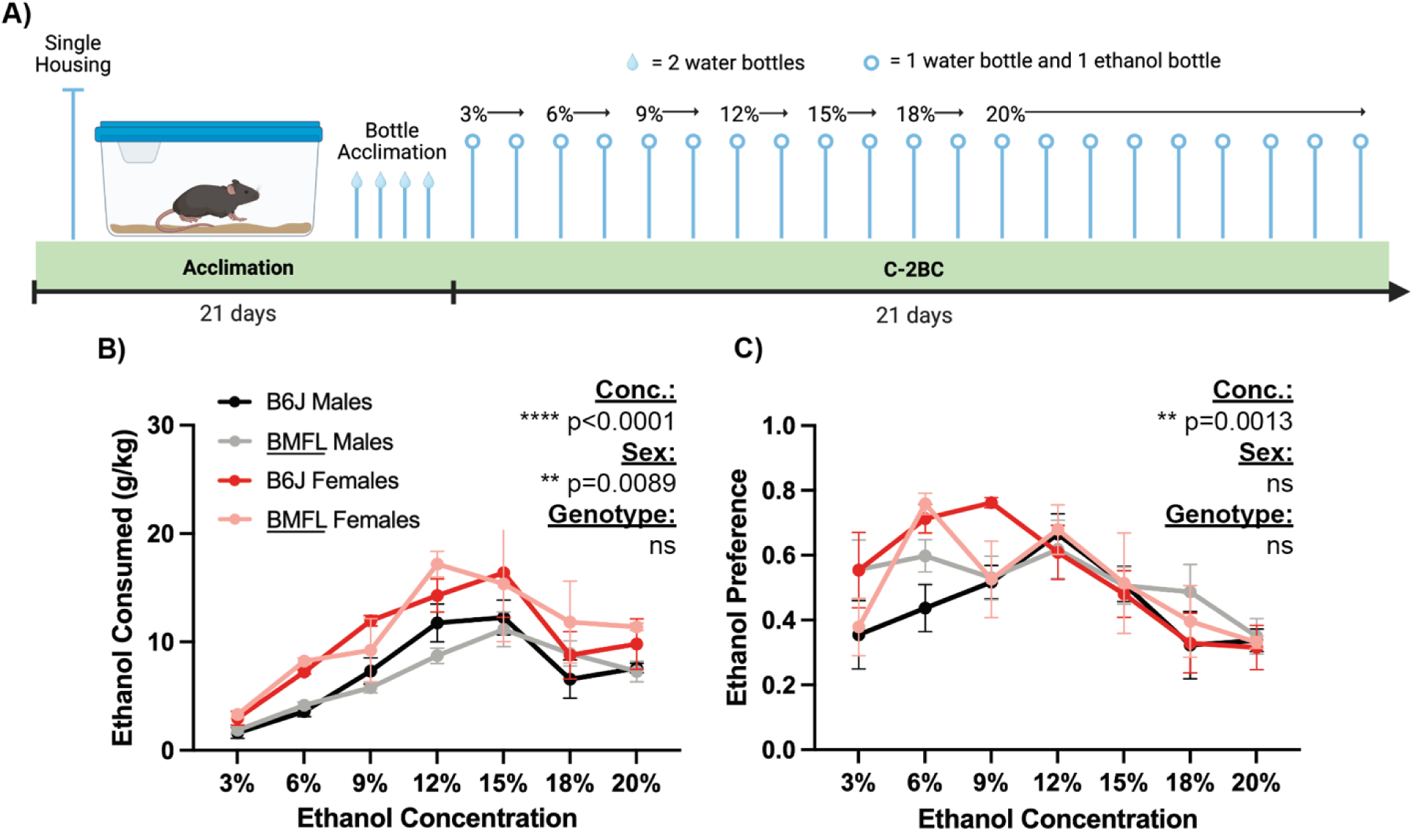
**(A)** Schematic detailing time course for single-housing, bottle acclimation, and continuous two-bottle choice experiments to assess impact of BMFL genotype on ethanol consumption (n=3 B6J males, n=3 B6J females, n=4 BMFL males, n=2 BMFL females). **(B)** There was no difference in ethanol consumed (g/kg) by BMFL mice versus B6J mice. A main effect of sex persists in both genotypes, with females consuming significantly more ethanol than males. **(C)** There is also no difference in ethanol preference in BMFL mice compared to B6J mice. Additionally there was no sex difference in ethanol preference in either genotype. #p<0.1, *p<0.05, **p<0.01, ***p<0.001, ****p<0.0001. Created in BioRender. Keefauver, T. (2026) https://BioRender.com/43o4vcr

### 3.3 Disrupting molecular clock function in NAc astrocytes controls binge-like drinking during drinking-in-the-light (DIL) and drinking-in-the-dark (DID) paradigms

Previously we demonstrated that mice with a loss of BMAL1 in NAc astrocytes self-administered more food pellets only in a progressive ratio task.^48^ Thus, we predicted models of binge-like ethanol consumption may more appropriately model this operant, motivated phenotype. DID is a validated model of binge-like ethanol consumption in the home cage, occurring early in the dark cycle when mouse ethanol consumption is highest.^53, 57^ We decided to additionally perform a similar but novel paradigm in the light cycle, referred to as drinking-in-the-light (DIL), so that we could measure diurnal differences in binge-like ethanol consumption.

We performed one cycle each of DIL and DID in the same cohort of virus injected BMFL mice to measure the effect of BMAL1 loss in NAc astrocytes on binge-like ethanol consumption (**Figure 5A**). There was no main effect of sex in DIL or DID (**Figure S2**), so sexes were collapsed for all figures and analyses. Loss of BMAL1 function in NAc astrocytes was associated with significant changes in ethanol consumption during both DIL and DID (**Figure 5: B-E**). During DIL, mice receiving the Cre virus consumed significantly more ethanol on days 1 and 4 only (**Figure 5B**) (<u>Virus:</u> F(1, 17) = 6.725, *p=0.0189; <u>Time:</u> F(2.113, 35.91) = 6.703, **p=0.0029; <u>Time x Virus:</u> F(2.113, 35.91) = 5.424, **p=0.0078; Sidak’s multiple comparisons: Day 1 **p_adj_=0.0023; Day 4 *p_adj_ =0.0269). There was no difference between virus groups in cumulative ethanol consumption throughout the DIL paradigm (**Figure 5C**). Mice did not show evidence of binge-like consumption during the 4-hour session of DIL on Day 4 (**Figure 5B**).

**Figure 5:**
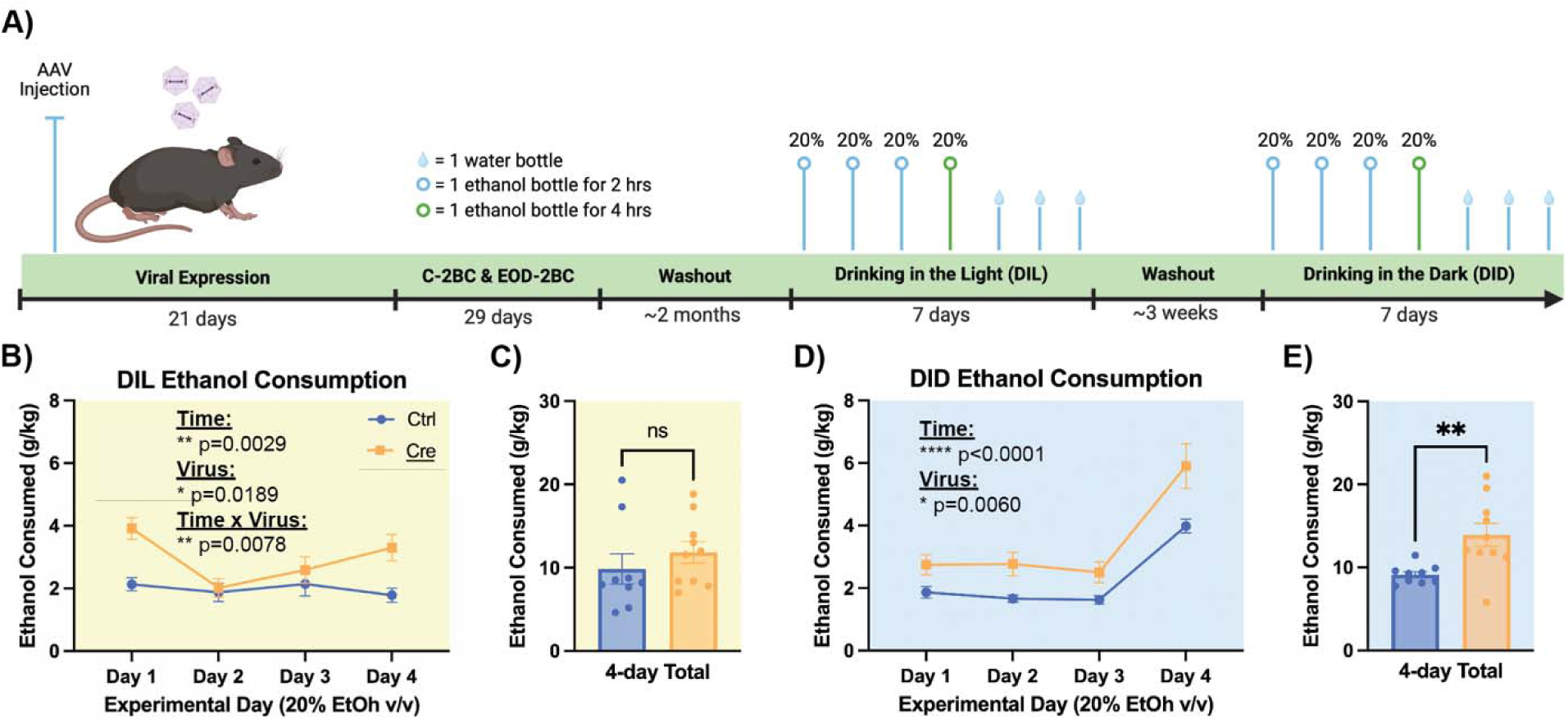
There was no main effect of sex in DIL or DID (**Figure S2**), so sexes were collapsed for all figures and analyses. **(A)** Schematic detailing time course before and during drinking-in-the-light (DIL) and drinking-in-the-dark (DID) experiments in BMFL mice (n=12 males; n=7 females). **(B)** Mice with a loss of BMAL1 function in NAc astrocytes (Gfap-Cre) consume significantly more alcohol compared to controls during DIL, and there is a significant interaction effect of time and virus. Mice receiving the Cre virus only consumed more ethanol on days 1 and 4 of the task, but show no evidence of binge drinking during the 4-hour session on Day 4 (green). **(C)** Cumulative ethanol consumption in the DIL task was not significantly different in mice with a loss of NAc astrocytic BMAL1. **(D)** Mice with a loss of BMAL1 function in NAc astrocytes consume significantly more alcohol compared to controls during DID. Additionally, all mice in this task show evidence of binge drinking during the 4-hour session on Day 4 (green). **(E)** Cumulative ethanol consumption in the DID task was 50% higher in mice with a loss of BMAL1 in NAc astrocytes. Experiments conducted in the following light conditions: Yellow = light cycle only; Blue = dark cycle only. #p<0.1, *p<0.05, **p<0.01, ***p<0.001, ****p<0.0001. Created in BioRender. Keefauver, T. (2026) https://BioRender.com/x0k76ct

During DID, mice receiving the Cre virus consumed significantly more ethanol throughout the entire paradigm (Virus: F(1, 17) = 9.825, **p=0.0060; <u>Time x Virus:</u> F(1.994, 33.90) = 1.506, p=ns, 0.2362; Sidak’s multiple comparisons: Day 1 #p_adj_=0.0790; Day 2 #p_adj_=0.0630; Day 3 #p_adj_=0.0790; Day 4 #p_adj_=0.0790) All mice showed significant evidence of binge-like consumption during the 4-hour session of DID on Day 4 (**Figure 5D**; <u>Time:</u> F(1.994, 33.90) = 46.98, ****p<0.0001; Sidak’s multiple comparisons: Day 4 vs. Days 1-3 **p_adj_<0.01; See **Supplementary Material** for individual Sidak’s multiple comparison test values). Cumulative ethanol consumption throughout the DID paradigm was also significantly different between virus groups (t(10.31) = 3.291, p=0.0078) with a large effect size (*d* =1.44) (**Figure 5E**).

### 3.4 Loss of BMAL1 function in NAc astrocytes is not associated with differences in locomotor response to novelty, sucrose preference, or social interaction

We next investigated whether differences in reward-related behaviors were associated with observed differences in ethanol consumption. The same cohort of virus injected BMFL mice was also tested for SPT, SIT, and LRN (**Figure 6A**). LRN is an established predictor of general drug-seeking propensity, and results were compared to alcohol consumption to parse potential alcohol-specific effects.^58, 59^ We also used LRN as a negative control to confirm mice with a loss of BMAL1 in NAc astrocytes show no changes in locomotion or behavioral rhythms, as previously established in the literature.^50^ SPT is a measure of anhedonic-like behavior,^60, 61^ allowing us to parse impacts of NAc astrocyte rhythm disruption on ethanol consumption compared to a natural reward. SIT is an established measure of preference for social novelty.^62^ We performed SIT to elucidate whether effects of NAc astrocyte rhythm disruption were specific to ethanol consumption or more generalized to novelty-seeking, including social novelty.

**Figure 6:**
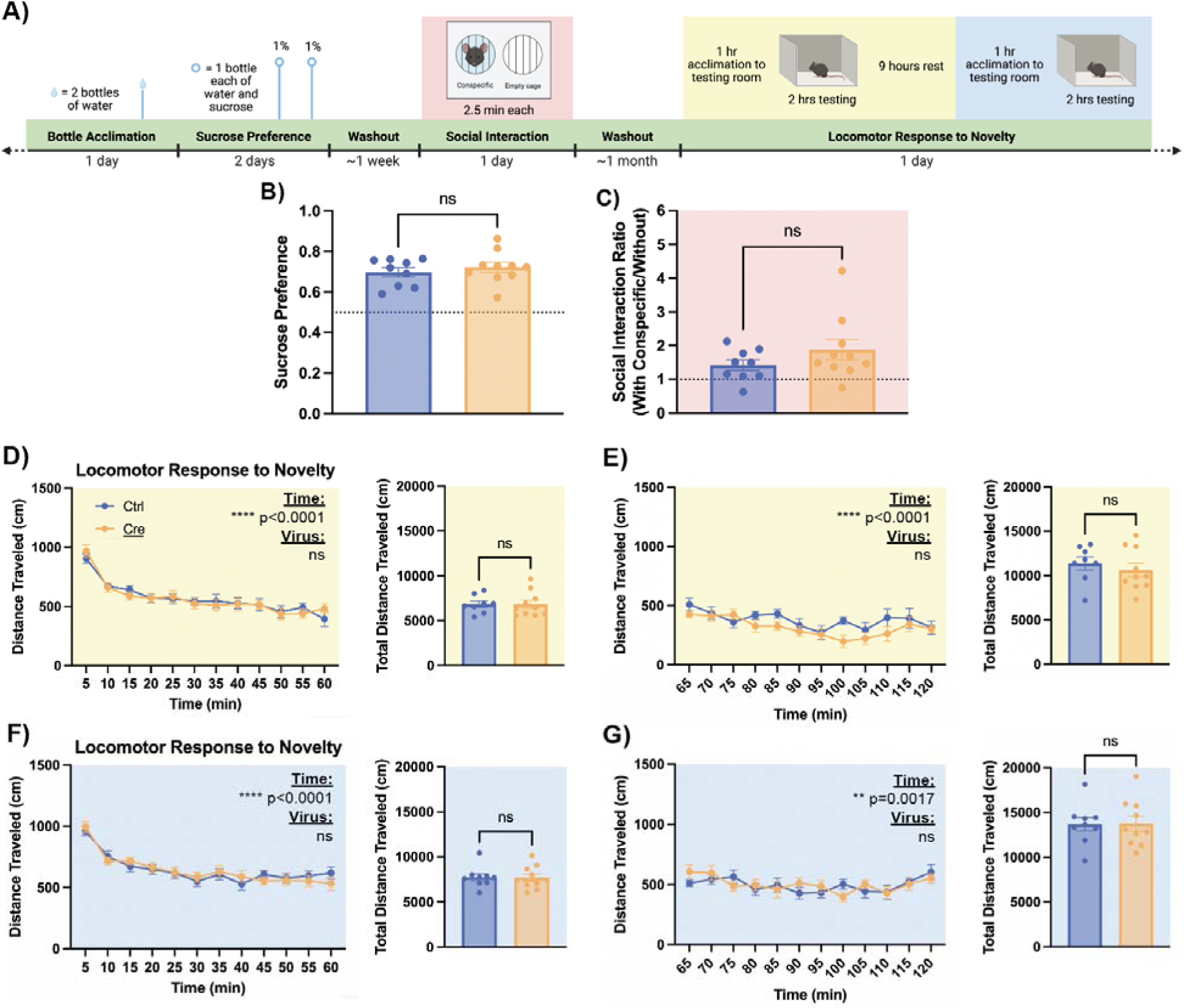
We found no main effect of sex in any of SPT, SIT, or LRN (**Figure S3**), so sexes were collapsed for all figures and analyses. **(A)** Schematic detailing time course for bottle acclimation sucrose preference test (SPT), social interaction test (SIT), and locomotor response to novelty (LRN) in BMFL mice (n=12 males; n=7 females). **(B)** Mice with loss of BMAL1 function in NAc astrocytes show no differences in sucrose preference compared to controls. **(C)** In a social interaction test performed with conspecifics, mice with a loss of BMAL1 function in NAc astrocytes show no difference in social interaction ratio (time spent with the conspecific/time spent without) compared to controls. **(D)** During the light cycle, mice with a loss of BMAL1 function in NAc astrocytes show no difference in LRN compared to controls. **(E)** There is no difference in habituation to LRN between ablated animals and controls during the light cycle. **(F)** During the dark cycle, mice with a loss of BMAL1 function in NAc astrocytes show no difference in LRN compared to controls. **(G)** There is also no difference in habituation to LRN between ablated animals and controls during the dark cycle. Experiments conducted in the following light conditions: Yellow = light cycle only; Blue = dark cycle only; Red = during the light cycle but under red light conditions. #p<0.1, *p<0.05, **p<0.01, ***p<0.001, ****p<0.0001. Created in BioRender. Keefauver, T. (2026) https://BioRender.com/z8o0u2s

We found no main effect of sex on SPT, SIT, or LRN (**Figure S3**), so sexes were collapsed for all figures and analyses. There were also no-order effects of assignment to light vs. dark first in LRN, so figures and analyses were collapsed. Sucrose preference was not impacted by loss of BMAL1 in NAc astrocytes (**Figure 6B**), and all mice preferred sucrose within a typical range for mice on a B6J background (∼60-80%).^63^ In SIT performed with conspecifics, loss of BMAL1 function in NAc astrocytes did not impact social interaction ratio (time spent with the conspecific/time spent without; **Figure 6C**). Loss of BMAL1 in NAc astrocytes did not impact LRN during the light cycle (**Figure 6D**) or dark cycle (**Figure 6F**). There was also no impact of BMAL1 functional ablation on within-session habituation to LRN (**Figure 6: E, G**).

## 4. Discussion

Circadian rhythm disruptions are key contributors to problematic alcohol use.^1–4^ While evidence suggests that alterations in circadian clock genes in NAc neurons impact ethanol-related behaviors,^6, 9^ no studies have examined astrocyte-specific effects. We hypothesized that alcohol consumption is controlled, at least in part, by astrocyte rhythms in the NAc. Astrocyte molecular clock function was disrupted in the NAc with viral injections of Gfap-Cre in mice with a floxed *Arntl* gene encoding for BMAL1, and mice were assessed for two-bottle choice drinking, binge-like drinking, locomotor response to novelty, sucrose preference, and social interaction. Disrupting molecular clock function in NAc astrocytes increased acute binge-like drinking during both the light cycle and dark cycle, but not 2BC drinking. Loss of BMAL1 function in NAc astrocytes was not associated with changes in reward-related behavior or locomotion. We also confirmed that mice used in this experiment, with a BMFL genotype, showed no difference in ethanol consumption or preference compared to B6J mice, the genotype typically used for voluntary drinking experiments.^64^ Results from LRN, SPT, and SIT show no differences between groups, suggesting genetic disruption of NAc astrocyte molecular clock function specifically impacts binge-like drinking, and that effects are not generalizable to novelty seeking or natural reward.

Previous research using chemogenetics identified a role for both NAc neurons^65^ and astrocytes^66^ in ethanol consumption, yet neither study explored molecular clock contributions. We demonstrate for the first time that disrupting NAc astrocyte molecular rhythms in NAc promotes increased ethanol consumption in a mouse model of binge-like drinking. Notably, disruption of astrocyte molecular rhythms selectively increased binge-like ethanol consumption without altering voluntary ethanol intake during 2BC. This specificity to binge-like drinking suggests astrocytic BMAL1 may preferentially regulate NAc processes engaged during episodes of acute, excessive ethanol consumption rather than baseline ethanol preference or general reward processing. Binge-like drinking paradigms are characterized by high levels of ethanol intake over a relatively short period of time^53, 57^ and are thought to rely heavily on NAc-mediated reward signaling and glutamatergic transmission.^67, 68^ In contrast, 2BC paradigms measure voluntary ethanol consumption across longer time scales and likely engage broader neural circuits involved in reinforcement learning, habit formation, and homeostatic regulation^69^ which may compensate for loss of astrocyte clock function over time. Our findings suggest astrocyte molecular rhythms may influence processing of ethanol intake during discrete drinking episodes rather than prolonged voluntary ethanol consumption.

Although the present study was not designed to identify molecular mechanisms underlying astrocyte clock regulation of alcohol drinking, previous work from our lab and others identified several clock-controlled astrocyte functions as plausible contributors.^48, 49, 70^ Astrocytes actively regulate extracellular glutamate concentrations, modulate GABAergic signaling, and provide metabolic support necessary for synaptic transmission.^68, 69^ Each of these processes exhibits circadian regulation and is influenced by BMAL1-dependent transcriptional programs.^70–78^ Disruption of astrocyte molecular rhythms could therefore alter temporal coordination of glutamate clearance or metabolic coupling within NAc,^79^ thus modifying neuronal activity during alcohol consumption. BMAL1 ablation in astrocytes is also associated with an overall increase in astrocyte reactivity,^80^ but it is unclear whether the markers of astrocyte reactivity [*e.g. Gfap* (cytoskeleton)*, Fabp7* (lipid handling)*, Mmp14* (extracellular matrix interactions)*, Cxcl5* (immune signaling)] are independent of clock-controlled pathways. Accordingly, behaviors observed in our study due to BMAL1 functional ablation may be attributable to alternative signaling pathways, in addition to clock-controlled mechanisms. Future mechanistic studies are necessary to determine which clock-controlled astrocyte function(s) mediates the relationship between astrocyte rhythms and binge-drinking.

Our findings should be interpreted in the larger context of a bidirectional relationship between circadian rhythm disruption and problematic alcohol use in humans.^5, 18, 19, 21^ The present study ablates the molecular clock in NAc astrocytes and assesses resulting changes in alcohol drinking. However, alcohol and alcohol misuse can also disrupt molecular clocks, leading to changes in cellular function or behavior.^81–84^ Clinical and preclinical studies consistently show disturbances in circadian rhythms,^20, 21^ including shift work,^22^ irregular sleep schedules,^3, 21^ and genetic variation in core clock genes,^10, 23, 24, 41^ are associated with increased risk for alcohol misuse.

While previous work has primarily focused on neuronal clock function,^6, 9^ the present study suggests astrocyte clock function also contributes to regulating alcohol consumption. Disruption of astrocyte molecular clocks may alter temporal organization of reward circuitry in ways that increase susceptibility to binge drinking. Our results expand understanding of how circadian dysfunction influences alcohol use disorder and highlight astrocytes as an important, understudied cellular target for future mechanistic and therapeutic investigations. Results from our study also provide astrocytic context to recent findings in NAc neurons.^9^ Using a similar viral-mediated *Cre* strategy, conditional knockout of BMAL1 in NAc neurons of BMFL mice was associated with an increase in voluntary ethanol consumption over 11 sessions of EOD-2BC in both males and females. While we did not observe a similar phenomenon in our EOD-2BC model for astrocytic BMAL1, we only administered 4 sessions of EOD-2BC. Future studies should repeat the astrocyte BMFL experiment and administer at least 10 sessions to determine whether effects of BMAL1 on intermittent ethanol consumption are cell-type specific in NAc.

The LRN findings from the present study contrast with a previously published LRN phenotype in BMFL mice with disrupted NAc astrocyte rhythmicity. Previous findings from BMFL mice receiving NAc injections of Gfap-Cre demonstrated that loss of BMAL1 in NAc astrocytes was associated with a light-cycle specific increase in total distance travelled during LRN.^48^ However, we did not observe this phenotype in the present study. Phenotypic differences are likely explained by two key differences in experimental design. First, the present study tested one group of animals during both the light and the dark (half tested in the light first and the other half tested in the dark first to control for order effects), whereas the previous study tested two cohorts, a light-cycle cohort and a dark-cycle cohort. Second, the present study maintained the mice in extended single-housing, whereas the previous study maintained the mice in group housing. Both experimental design choices likely reduced the novelty of the LRN apparatus and task to the mice in the present study and therefore may have contributed to our lack of an observed light-cycle specific phenotype. The differences might also be explained by differences in the age of the mice. Mice in our previous experiment were 10-14 weeks old at the time of testing, whereas mice in our current experiment were 31-34 weeks during LRN. Future experiments should continue to assess mice with LRN and assist with further phenotyping of BMFL mice with cell-type specific manipulations.

Our study has four main limitations. First, we did not collect mouse blood ethanol concentrations (BECs) during binge-like drinking assays (DIL and DID). Since BMFL and B6J animals consumed identical but low amounts of ethanol, we expect BECs for our animals would have fallen on the low end of reported ranges from numerous B6J DID experiments (∼0.8-1.6mg/mL).^53, 57, 85^ In future studies, we will repeat our binge-like drinking experiments in BMFL mice and will employ technical improvements: we will record BECs after each session using tail bleeds and we will administer multiple weeks each of DIL and DID to measure escalation of ethanol consumption over time.^86^ Second, each behavioral experiment is limited by possible carry-over effects from previous exposure to ethanol, previous behavioral testing, and extended time spent in single-housing due to experimental design. While extensive 3-4 week washout periods were implemented to minimize carry-over effects, it is possible that mice in subsequent drinking experiments drank less ethanol because they were ethanol-naïve. It is also possible that our binge-like drinking results would not replicate in ethanol-naïve mice. Future studies should repeat each drinking experiment with a separate cohort of mice to determine if carry-over effects or extended single-housing impact results. Third, we did not measure ethanol clearance, metabolism, or bitter tastant (*e.g.,* quinine) perception^87^ in any of our drinking experiments. This limits the interpretability of our findings, especially given that we observe relatively low ethanol consumption (g/kg) in all genotypes of our mice. Future drinking experiments will measure each of these variables to determine whether our mice metabolize and clear ethanol appropriately and whether bitter taste perception impacts ethanol preference in our animals. Fourth, results are limited by a moderate sample size. Our future experiments will use our own results alongside recent literature to conduct power analyses, determine effect size, and design new surgical cohorts with high statistical power to detect differences during diurnal drinking experiments.

## 5. Conclusions

Our study provides the first evidence for a role of NAc astrocyte rhythmicity in binge-like alcohol consumption, carving out a unique role for astrocytes in controlling temporal organization of reward circuitry and susceptibility to binge-drinking. Future studies will investigate clock-controlled astrocyte mechanisms, such as extrasynaptic glutamate uptake and ATP release,^46, 47^ that may underlie binge-like drinking behavior. Understanding bidirectional relationships between astrocyte molecular rhythms and alcohol consumption will inform development of astrocyte-specific therapeutics to treat circadian symptoms and risk factors of problematic alcohol use.

## Acknowledgements

We thank Drs. Max Joffe and Colleen McClung for helpful discussions and written comments. We thank Micah Shelton, Dr. Kaitlyn Petersen, and Dr. Taylor Stowe for technical assistance and helpful discussions. We thank the University of Pittsburgh Department of Psychiatry and Dr. David Lewis for material support. This research was supported by the National Institute on Alcohol Abuse and Alcoholism [NIAAA R21 (AA031074; MPIs: KDK, MLS, RWL; Co-Is: SPF, GEH)], the National Institute of Mental Health [NIMH K01 (MH128763; KDK)], and the National Institute of Neurological Disorders and Stroke [predoctoral training grant to TK (NINDS: T32 NS141747)].

## Data Availability Statement

The behavioral data supporting the findings of this study are openly available on GitHub repository at https://github.com/torikeefauver/CircadianAstrocyteAlcoholPaper. The imaging data supporting the findings of this study are available from the corresponding authors upon reasonable request, due to large file sizes.

## Ethics Approval Statement

Ethical approval for the use of vertebrate animals in this study was obtained on 4/18/2024 from the University of Pittsburgh Institutional Animal Care and Use Committee (IACUC), protocol #: 24044835.

