## Supplementary Material for "Astrocyte molecular rhythm disruption in nucleus accumbens promotes increased binge-like drinking in mice"

**Contents**

I. **Supplementary Figures**
II. **Extended figure captions including full statistics**

**I. Supplementary Figures**

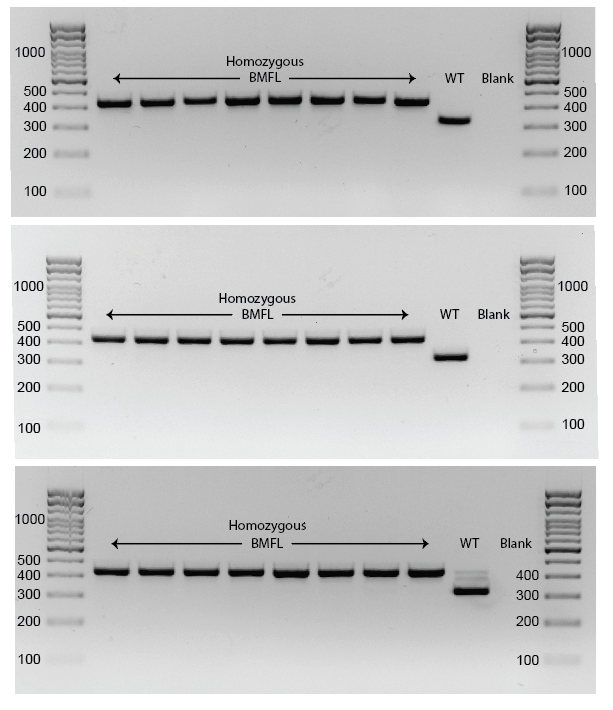

Figure S1: Homozygous genotype confirmation for BMFL mice.

Homozygous genotypes of BMFL mice were confirmed using PCR and gel electrophoresis (n=12 males; n=7 females). Primers suggested by the JAX website were used (JAX:007668; Forward primer: oIMR7525 = ACT GGA AGT AAC TTT ATC AAA CTG; Reverse primer: oIMR7526 = CTG ACC AAC TTG CTA ACA ATT A). Expected band sizes: Homozygous foxed = 431bp, Heterozygous floxed = 431bp and 327bp, Wild type = 327 bp.

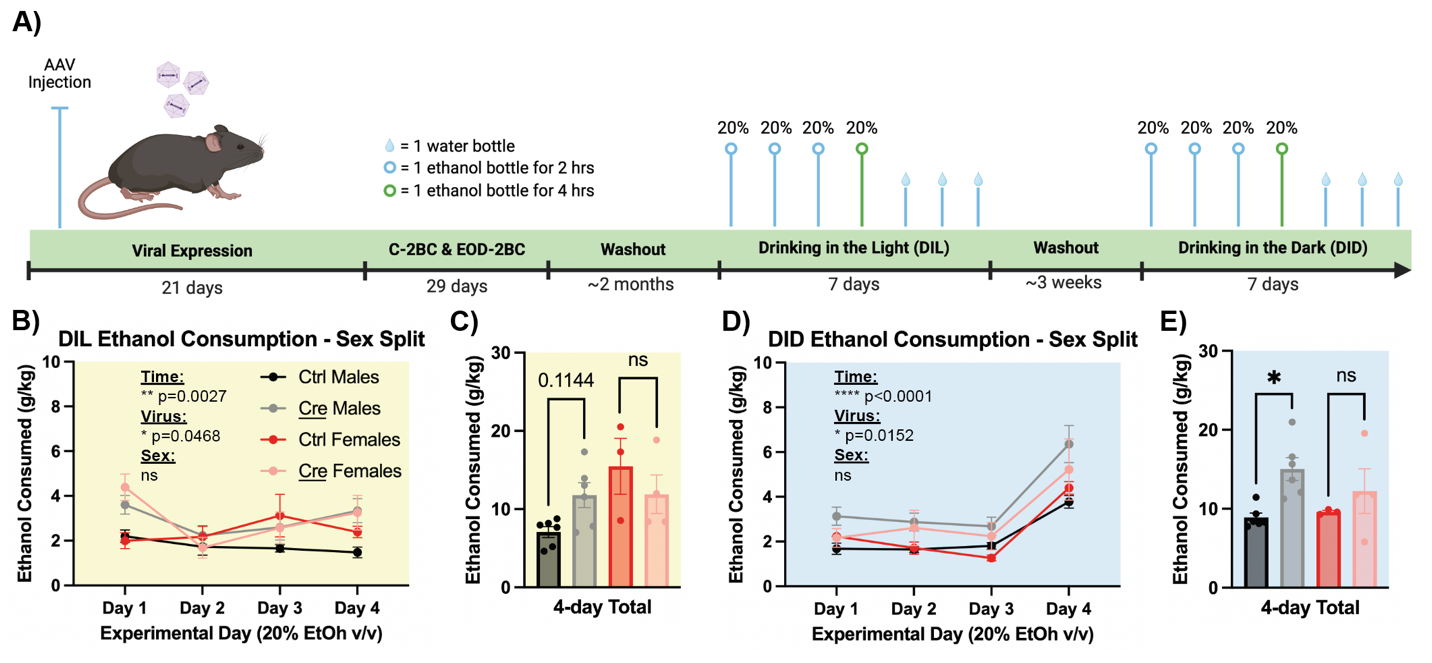

Figure S2: Sex-split figures for DIL/DID (complementary to Figure 5)

**(A)** Schematic detailing time course before and during drinking-in-the-light (DIL) and drinking-in-the-dark (DID) experiments in BMFL mice (n=12 males; n=7 females). **(B)** Mice with a loss of BMAL1 function in NAc astrocytes (Gfap-Cre) consume significantly more alcohol compared to controls during DIL, and there is a significant interaction effect of time and virus. We found no main effect of sex on one cycle of DIL ethanol consumption, so sex was collapsed in the main manuscript (Figure 5). **(C)** Cumulative ethanol consumption in the DIL task was not significantly different in mice with a loss of NAc astrocytic BMAL1, but there was a significant difference in cumulative ethanol consumption by sex, with females consuming more ethanol concurrent with established literature.^1, 2^ Within-sex comparisons reveal no effect of virus in females, and a near-trend of an effect of virus in males. **(D)** During DID, mice with a loss of BMAL1 function in NAc astrocytes consume significantly more alcohol compared to controls. We found no main effect of sex on one cycle of DID ethanol consumption, so sex was collapsed in the main manuscript (Figure 5). **(E)** Cumulative ethanol consumption in the DID task was significantly higher in mice with a loss of BMAL1 in NAc astrocytes, and there is no main effect of sex. Within-sex comparisons reveal no effect of virus in females, and a significant effect of virus in males. Experiments conducted in the following light conditions: Yellow = light cycle only; Blue = dark cycle only. #p<0.1, *p<0.05, **p<0.01, ***p<0.001, ****p<0.0001.

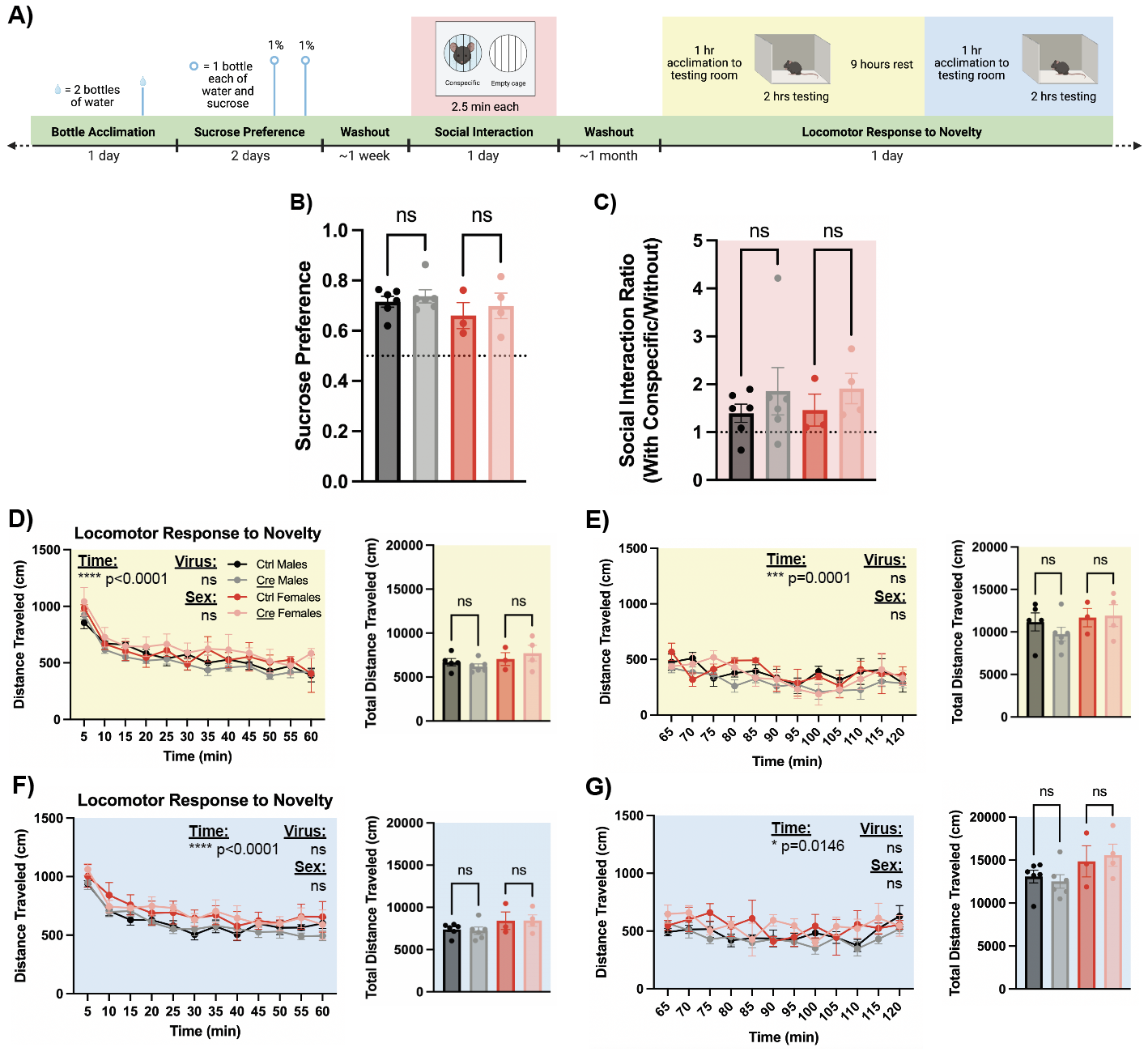

Figure S3: Sex-split figures for behavior (complementary to Figure 6)

**(A)** Schematic detailing time course for bottle acclimation, sucrose preference test (SPT), social interaction test (SIT), and locomotor response to novelty (LRN) in BMFL mice (n=12 males; n=7 females). **(B)** Mice with loss of BMAL1 function in NAc astrocytes show no differences in sucrose preference compared to controls or between sexes. Within-sex comparisons reveal no effect of virus in males or females. **(C)** In a social interaction test performed with conspecifics, mice with a loss of BMAL1 function in NAc astrocytes show no difference in social interaction ratio (time spent with the conspecific/time spent without) compared to controls or between sexes. Within-sex comparisons reveal no effect of virus in males or females. **(D)** During the light cycle, mice with a loss of BMAL1 function in NAc astrocytes show no difference in LRN compared to controls, and there is no main effect of sex. Within-sex comparisons reveal no effect of virus in males or females. **(E)** There is no difference in habituation to LRN between ablated animals and controls or between sexes during the light cycle. Within-sex comparisons reveal no effect of virus in males or females. **(F)** During the dark cycle, mice with a loss of BMAL1 function in NAc astrocytes show no difference in LRN compared to controls and there is a trend towards a main effect of sex. Within-sex comparisons reveal no effect of virus in males or females. **(G)** There is also no difference in habituation to LRN between ablated animals and controls or between sexes during the dark cycle or between sexes during the dark cycle. Within-sex comparisons reveal no effect of virus in males or females. Experiments conducted in the following light conditions: Yellow = light cycle only; Blue = dark cycle only; Red = during the light cycle but under red light conditions. #p<0.1, *p<0.05, **p<0.01, ***p<0.001, ****p<0.0001.

**II. Extended figure captions including full statistics:**

Figure 1 Extended Caption:

**(A)** Schematic illustrating bilateral stereotaxic injections of either AAV8-Gfap-Cre-GFP or AAV8-Gfap-eGFP virus into NAc of BMFL mice; injections at 1uL/min, coordinates relative to bregma: AP: +1.5, ML: ± 1.5, and DV: −4.4; angle 10°. AC=Anterior Commissure. **(B)** Immunohistochemistry (IHC) images confirm Gfap-Cre causes loss of BMAL1. **(C)** Images also confirm astrocyte-specific tropism of AAV8 viruses. Created in BioRender. Keefauver, T. (2026) <https://BioRender.com/fli9rh2>

Figure 2 Extended Caption:

Schematic illustrating full experimental timeline for BMFL mice (n=12 males; n=7 females) receiving viral injections. Experiments conducted in the following light conditions: Yellow = light cycle only; Blue = dark cycle only; Yellow/Blue gradient = 24/7 but with separate measurements in light and dark cycles; Red = during the light cycle but under red light conditions. Created in BioRender. Keefauver, T. (2026) <https://BioRender.com/r82ixtj>

Figure 3 Extended Caption:

**(A)** Schematic detailing time course for viral injections, bottle acclimation, continuous two-bottle choice (C-2BC), and every other day two-bottle choice (EOD-2BC) experiments in BMFL mice (n=12 males; n=7 females). **(B)** Mice with a loss of BMAL1 function in NAc astrocytes (Gfap-Cre) show no differences in ethanol consumption (g/kg) in the light cycle during C-2BC drinking, and females consumed significantly more alcohol than males (Concentration: F(2.466, 36.98) = 10.55, **** p<0.0001; Sex: F(1, 15) = 6.605, *p=0.0213; Virus: F(1, 15) = 1.058, p=0.32; Concentration × Sex: F(2.466, 36.98) = 1.017, p=0.3847; Concentration x Virus: F(2.466, 36.98) = 0.6170, p=0.5777; Sex x Virus: F(1, 15) = 2.774, p=0.1165; Concentration x Sex x Virus: F(2.466, 36.98) = 0.5126, p=0.6409). **(C)** During the dark cycle, females still consume more ethanol than males, and there is a trend towards a significant effect of BMAL1 loss on ethanol consumption (Concentration: F(3.796, 56.31) = 16.37, ****p<0.0001; Sex: F(1, 15) = 20.14, ***p=0.0004; Virus: F(1, 15) = 4.342, #p=0.0547; Concentration × Sex: F(3.796, 56.31) = 1.445, p=0.2231; Concentration x Virus: F(3.796, 56.31) = 0.5811, p=0.6689; Sex x Virus: F(1, 15) = 0.09456, p=0.7627; Concentration x Sex x Virus: F(3.796, 56.31) = 0.9534, p=0.4369). **(D)** When data is collapsed into 24-hour periods, results mimic the dark cycle (Concentration: F(6, 88) = 20.06, ****p<0.0001; Sex: F(1, 15) = 17.69, ***p=0.0008; Virus: F(1, 15) = 3.871, #p=0.0679; Concentration × Sex: F(6, 88) = 0.8657, p=0.5235; Concentration x Virus: F(6, 88) = 1.216, p=0.3058; Sex x Virus: F(1, 15) = 0.9801, p=0.3379; Concentration x Sex x Virus: F(6, 88) = 1.161, p=0.3347). **(E)** Mice with a loss of BMAL1 function in NAc astrocytes show no differences in ethanol preference in the light cycle during C-2BC drinking, and females preferred ethanol significantly more than males (Concentration: F(2.948, 44.23) = 2.487, #p=0.0738; Sex: F(1, 15) = 4.637, *p=0.0480; Virus: F(1, 15) = 0.01144, p=0.9163; Concentration × Sex: F(2.948, 44.23) = 0.9197, p=0.4378; Concentration x Virus: F(2.948, 44.23) = 1.126, p=0.3483; Sex x Virus: F(1, 15) = 1.976, p=0.1802; Concentration x Sex x Virus: F(2.948, 44.23) = 1.082, p=0.3660). **(F)** During the dark cycle, there is a significant effect of ethanol concentration on ethanol preference and a trend towards a significant effect of sex (Concentration: F(6, 90) = 19.54, ****p<0.0001; Sex: F(1, 15) = 3.313, #p=0.0887; Virus: F(1, 15) = 1.890, p=0.1894; Concentration × Sex: F(6, 90) = 1.370, p=0.2351; Concentration x Virus: F(6, 90) = 1.230, p=0.2987; Sex x Virus: F(1, 15) = 1.164, p=0.2976; Concentration x Sex x Virus: F(6, 90) = 1.230, p=0.2986). **(G)** When data is collapsed into 24-hour periods, results mimic the dark cycle (Concentration: F(4.107, 61.60) = 16.54, ****p<0.0001; Sex: F(1, 15) = 4.393, #p=0.0535; Virus: F(1, 15) = 2.172, p=0.1612; Concentration × Sex: F(4.107, 61.60) = 1.219, p=0.3119; Concentration x Virus: F(4.107, 61.60) = 1.737, p=0.1517; Sex x Virus: F(1, 15) = 1.663, p=0.2168; Concentration x Sex x Virus: F(4.107, 61.60) = 1.379, p=0.2508) **(H)** Mice with loss of BMAL1 function in astrocytes show no within-sex differences in cumulative ethanol consumption across the 21-day task (Sidak’s multiple comparisons; Ctrl Males vs. Cre Males: p_adj_ = 0.3588; Ctrl Females vs. Cre Females: p_adj_ = 0.8001). There is no main effect of virus on cumulative ethanol consumption, but there is a main effect of sex on cumulative ethanol consumption (Virus: F(1,15) = 1.675, p=0.2152**;** Sex: F(1, 15) = 15.62, **p=0.0013**). (I)** In an EOD-2BC drinking task, mice show main effects only of sex and time on ethanol consumption, with females consuming more ethanol than males, and both sexes increasing their drinking across the 8-days (Time: F(1.986, 28.47) = 8.161, **p=0.0016; Sex: F(1, 15) = 13.03, **p=0.0026; Virus: F(1, 15) = 0.07164, p=0.7926; Time x Sex: F(1.986, 28.47) = 0.7594, p=0.4763; Time x Virus: F(1.986, 28.47) = 0.7706, p=0.4713; Sex x Virus: F(1, 15) = 0.1991, p=0.6618; Time x Sex x Virus: F(1.986, 28.47) = 0.4895, p=0.6167). **(J)** Similar to EOD-2BC ethanol consumption, there is a significant effect of time and a trend towards a significant effect of sex on ethanol preference. There is an additional significant interaction between time and sex, with only females exhibiting a stronger preference for ethanol at the end of the task (Time: F(2.665, 51.52) = 13.41, ****p<0.0001; Sex: F(1, 20) = 3.816, #p=0.0649; Virus: F(1, 20) = 0.01253, p=0.9120; Time x Sex: F(2.665, 51.52) = 2.909, p=0.0491; Time x Virus: F(2.665, 51.52) = 0.1027, p=0.9448; Sex x Virus: F(1, 20) = 0.01033, p=0.9201; Time x Sex x Virus: F(2.665, 51.52) = 0.02095, p=0.9929). Experiments conducted in the following light conditions: Yellow = light cycle only; Blue = dark cycle only. #p<0.1, *p<0.05, **p<0.01, ***p<0.001, ****p<0.0001. Created in BioRender. Keefauver, T. (2026) <https://BioRender.com/f4ibihx>

Figure 4 Extended Caption:

**(A)** Schematic detailing time course for single-housing, bottle acclimation, and continuous two-bottle choice experiments to assess impact of BMFL genotype on ethanol consumption (n=3 B6J males, n=3 B6J females, n=4 BMFL males, n=2 BMFL females). **(B)** There is no difference in ethanol consumed (g/kg) by BMFL mice versus B6J mice. A main effect of sex persists in both genotypes, with females consuming significantly more ethanol than males (Concentration: F(2.586, 20.69) = 32.94, ****p<0.0001; Sex: F(1, 8) = 11.81, **p=0.0089; Genotype: F(1, 8) = 0.02698, p=0.8736; Concentration x Sex: F(2.586, 20.69) = 0.9119, p=0.4398; Concentration x Genotype: F(2.586, 20.69) = 1.247, p=0.3151; Sex x Genotype: F(1, 8) = 0.3082, p=0.5940; Concentration x Sex x Genotype: F(2.586, 20.69) = 0.7130, p=0.5358). **(C)** There is also no difference in ethanol preference in BMFL mice compared to B6J mice. Additionally there is no sex difference in ethanol preference in either genotype (Concentration: F(2.300, 18.02) = 9.085, **p=0.0013; Sex: F(1, 8) = 1.236, p=0.2985; Genotype: F(1, 8) = 0.4181, p=0.5360; Concentration x Sex: F(2.300, 18.02) = 1.360, p=0.2841; Concentration x Genotype: F(2.300, 18.02) = 0.8911, p=0.4407; Sex x Genotype: F(1, 8) = 1.845, p=0.2115; Concentration x Sex x Genotype: F(2.300, 18.02) = 1.184, p=0.3346). #p<0.1, *p<0.05, **p<0.01, ***p<0.001, ****p<0.0001. Created in BioRender. Keefauver, T. (2026) <https://BioRender.com/43o4vcr>

Figure 5 Extended Caption:

There was no main effect of sex in DIL or DID (**Figure S2**), so sexes were collapsed for all figures and analyses. **(A)** Schematic detailing time course before and during drinking-in-the-light (DIL) and drinking-in-the-dark (DID) experiments in BMFL mice (n=12 males; n=7 females). **(B)** Mice with a loss of BMAL1 function in NAc astrocytes (Gfap-Cre) consume significantly more alcohol compared to controls during DIL (Virus: F(1, 17) = 6.725, *p=0.0189), and there is a significant interaction effect of time and virus (Time: F(2.113, 35.91) = 6.703, **p=0.0029; Time x Virus: F(2.113, 35.91) = 5.424, **p=0.0078). Mice receiving the Cre virus only consumed more ethanol on days 1 and 4 of the task (Sidak’s multiple comparisons; Day 1 **p_adj_=0.0023; Day 4 *p_adj_ =0.0269), but show no evidence of binge drinking during the 4-hour session on Day 4 (green). We found no main effect of sex on one cycle of DIL ethanol consumption, so sex was collapsed (Sex: F(1, 15) = 0.7253, p=0.4078). **(C)** Cumulative ethanol consumption in the DIL task was not significantly different in mice with a loss of NAc astrocytic BMAL1 (t(14.82) = 0.8906, p=0.3874, d=0.4163). **(D)** Mice with a loss of BMAL1 function in NAc astrocytes consume significantly more alcohol compared to controls during DID (Virus: F(1, 17) = 9.825, **p=0.0060; Time x Virus: F(1.994, 33.90) = 1.506, p=ns, 0.2362; Sidak’s multiple comparisons: Day 1 #p_adj_=0.0790; Day 2 #p_adj_=0.0630; Day 3 #p_adj_=0.0790; Day 4 #p_adj_=0.0790). Additionally, all mice in this task show evidence of binge drinking during the 4-hour session on Day 4 (green) (Time: F(1.994, 33.90) = 46.98, ****p<0.0001; Sidak’s multiple comparisons: Day 4 vs. Days 1-3 **p_adj_<0.01; Ctrl_Day4_ vs. Ctrl_Day1_ ****p_adj_<0.0001; Ctrl_Day4_ vs. Ctrl_Day2_ ****p_adj_<0.0001; Ctrl_Day4_ vs. Ctrl_Day3_ ****p_adj_<0.0001; Cre_Day4_ vs. Cre_Day1_ **p_adj_=0.0018; Cre_Day4_ vs. Cre_Day2_ ***p_adj_=0.0008; Cre_Day4_ vs. Cre_Day3_ ***p_adj_=0.0002). We found no main effect of sex on one cycle of DID ethanol consumption, so sex was collapsed (Sex: F(1, 15) = 0.4346, p=0.5197). **(E)** Cumulative ethanol consumption in the DID task was 50% higher in mice with a loss of BMAL1 in NAc astrocytes (t(10.31) = 3.291, p=0.0078, d=1.44). Experiments conducted in the following light conditions: Yellow = light cycle only; Blue = dark cycle only. #p<0.1, *p<0.05, **p<0.01, ***p<0.001, ****p<0.0001. Created in BioRender. Keefauver, T. (2026) <https://BioRender.com/x0k76ct>

Figure 6 Extended Caption:

We found no main effect of sex in any of SPT, SIT, or LRN (**Figure S3**), so sexes were collapsed for all figures and analyses. **(A)** Schematic detailing time course for bottle acclimation, sucrose preference test (SPT), social interaction test (SIT), and locomotor response to novelty (LRN) in BMFL mice (n=12 males; n=7 females). **(B)** During the light cycle, mice with a loss of BMAL1 function in NAc astrocytes show no difference in LRN compared to controls (Time: F(6.093, 97.48) = 38.92, ****p<0.0001; Virus: F(1,16) = 0.0004492, p=0.9834; Time x Virus: F(6.093, 97.48) = 0.9873, p=0.4389; t(15.86) = 0.02201, p=0.9827, d=0.009731). **(C)** There is no difference in habituation to LRN between ablated animals and controls during the light cycle (Time: F(6.731, 107.7) = 5.262, ****p<0.0001; Virus: F(1,16) = 1.679, p=0.2135; Time x Virus: F(6.731, 107.7) = 0.1.237, p=0.2900; t(15.84) = 0.7165, p=0.4841, d=0.3342). **(D)** During the dark cycle, mice with a loss of BMAL1 function in NAc astrocytes show no difference in LRN compared to controls (Time: F(5.431, 92.33) = 40.53, ****p<0.0001; Virus: F(1,17) = 0.001513, p=0.9694; Time x Virus: F(5.431, 92.33) = 1.425, p=0.2185; t(16.97) = 0.03906, p=0.9693, d=0.01797). **(E)** There is also no difference in habituation to LRN between ablated animals and controls during the dark cycle (Time: F(5.959, 101.3) = 3.843, **p=0.0017; Virus: F(1,17) = 0.01725, p=0.8971; Time x Virus: F(5.959, 101.3) = 1.726, p=0.1231; t(16.99) = 0.05182, p=0.9593, d=0.02356). **(F)** Mice with loss of BMAL1 function in NAc astrocytes show no differences in sucrose preference compared to controls (t(16.98) = 0.7327, p=0.4737, d=0.3342). **(G)** In a social interaction test performed with conspecifics, mice with a loss of BMAL1 function in NAc astrocytes show no difference in social interaction ratio (time spent with the conspecific/time spent without) compared to controls (t(13.27) = 1.333, p=0.2051, d=0.5591). We found no main effect of sex on any of SPT, SIT, or LRN behaviors, so sex was collapsed throughout analyses (Sex_LRN, Light_: F(1, 14) = 2.673, p= 0.1243; Sex_LRN, Dark_: F(1,15) = 4.052, #p=0.0624; Sex_SPT_: F(1,15) = 1.741, p=0.2069; Sex_SIT_: F(1,15) = 0.02151, p=0.8853). Experiments conducted in the following light conditions: Yellow = light cycle only; Blue = dark cycle only; Red = during the light cycle but under red light conditions. #p<0.1, *p<0.05, **p<0.01, ***p<0.001, ****p<0.0001. Created in BioRender. Keefauver, T. (2026) <https://BioRender.com/z8o0u2s>

Figure S1: Homozygous genotype confirmation for BMFL mice.

Homozygous genotypes of BMFL mice were confirmed using PCR and gel electrophoresis (n=12 males; n=7 females). Primers suggested by the JAX website were used (JAX:007668; Forward primer: oIMR7525 = ACT GGA AGT AAC TTT ATC AAA CTG; Reverse primer: oIMR7526 = CTG ACC AAC TTG CTA ACA ATT A). Expected band sizes: Homozygous foxed = 431bp, Heterozygous floxed = 431bp and 327bp, Wild type = 327 bp.

Figure S2: Sex-split figures for DIL/DID (complementary to Figure 5)

**(A)** Schematic detailing time course before and during drinking-in-the-light (DIL) and drinking-in-the-dark (DID) experiments in BMFL mice (n=12 males; n=7 females). **(B)** Mice with a loss of BMAL1 function in NAc astrocytes (Gfap-Cre) consume significantly more alcohol compared to controls during DIL (Virus: F(1, 15) = 4.693, *p=0.0468), and there is a significant interaction effect of time and virus (Time: F(2.191, 32.86) = 6.767, **p=0.0027; Time x Virus: F(2.191, 32.86) = 6.924, **p=0.0024). We found no main effect of sex on one cycle of DIL ethanol consumption, so sex was collapsed in the main manuscript (Figure 5) (Sex: F(1, 15) = 0.7253, p=0.4078). **(C)** Cumulative ethanol consumption in the DIL task was not significantly different in mice with a loss of NAc astrocytic BMAL1 (Virus: F(1, 15) = 0.09156, p=0.7664), but there was a significant difference in cumulative ethanol consumption by sex, with females consuming more ethanol concurrent with established literature (Sex: F(1,15) = 4.934, *p=0.0421). Within-sex comparisons reveal no effect of virus in females (Sidak’s multiple comparisons; **p_adj_=0.4552), and a near-trend of an effect of virus in males (Sidak’s multiple comparisons; p_adj_=0.1144). **(D)** During DID, mice with a loss of BMAL1 function in NAc astrocytes consume significantly more alcohol compared to controls (Virus: F(1, 15) = 7.512, *p=0.0152). We found no main effect of sex on one cycle of DID ethanol consumption, so sex was collapsed in the main manuscript (Figure 5) (Sex: F(1, 15) = 0.4346, p=0.5197). **(E)** Cumulative ethanol consumption in the DID task was significantly higher in mice with a loss of BMAL1 in NAc astrocytes (Virus: F(1,15) = 7.512, *p=0.0152), and there is no main effect of sex (Sex: F(1,15) = 0.4346, p=0.5197). Within-sex comparisons reveal no effect of virus in females (Sidak’s multiple comparisons; p_adj_=0.5367), and a significant effect of virus in males (Sidak’s multiple comparisons; *p_adj_=0.0123). Experiments conducted in the following light conditions: Yellow = light cycle only; Blue = dark cycle only. #p<0.1, *p<0.05, **p<0.01, ***p<0.001, ****p<0.0001. Created in BioRender. Keefauver, T. (2026) <https://BioRender.com/x0k76ct>

Figure S3: Sex-split figures for behavior (complementary to Figure 6)

**(A)** Schematic detailing time course for bottle acclimation, sucrose preference test (SPT), social interaction test (SIT), and locomotor response to novelty (LRN) in BMFL mice (n=12 males; n=7 females). **(B)** Mice with loss of BMAL1 function in NAc astrocytes show no differences in sucrose preference compared to controls (Virus: F(1, 15) = 0.7136, p=0.4115) or between sexes (Sex: F(1. 15) = 1.741, p=0.2069). Within-sex comparisons reveal no effect of virus in males (Sidak’s multiple comparisons; **p_adj_=0.8586) or females (Sidak’s multiple comparisons; p_adj_=0.7564). **(C)** In a social interaction test performed with conspecifics, mice with a loss of BMAL1 function in NAc astrocytes show no difference in social interaction ratio (time spent with the conspecific/time spent without) compared to controls (Virus: F(1, 15) = 1.317, p=0.2690) or between sexes (Sex: F(1, 15) = 0.02151, p=0.8853). Within-sex comparisons reveal no effect of virus in males (Sidak’s multiple comparisons; **p_adj_=0.5781) or females (Sidak’s multiple comparisons; p_adj_=0.7383). **(D)** During the light cycle, mice with a loss of BMAL1 function in NAc astrocytes show no difference in LRN compared to controls (Time: F(5.668, 79.35) = 35.86, ****p<0.0001; Virus: F(1, 14) = 0.02797, p=0.8696), and there is no main effect of sex (Sex: F(1, 14) = 2.673, p=0.1243). Within-sex comparisons reveal no effect of virus in males (Sidak’s multiple comparisons; **p_adj_=0.7524) or females (Sidak’s multiple comparisons; p_adj_=0.7087). **(E)** There is no difference in habituation to LRN between ablated animals and controls (Time: F(6.082, 85.15) = 5.260, ***p=0.0001; Virus: F(1, 14) = 1.283, p=0.2763; Virus_Total_: F(1, 14) = 0.3053, p=0.5893) or between sexes during the light cycle (Sex: F(1, 14) = 0.4752, p=0.5019; Sex_Total_: F(1, 14) = 1.541, p=0.2348). Within-sex comparisons reveal no effect of virus in males (Sidak’s multiple comparisons; **p_adj_=0.5168) or females (Sidak’s multiple comparisons; p_adj_=0.9882). **(F)** During the dark cycle, mice with a loss of BMAL1 function in NAc astrocytes show no difference in LRN compared to controls (Time: F(5.069, 76.03) = 35.53, ****p<0.0001; Virus: F(1, 15) = 0.01749, p=0.8966) and there is a trend towards a main effect of sex (Sex: F(1, 15) = 4.052, #p=0.0624). Within-sex comparisons reveal no effect of virus in males (Sidak’s multiple comparisons; **p_adj_=0.9655) or females (Sidak’s multiple comparisons; p_adj_=0.9999). **(G)** There is also no difference in habituation to LRN between ablated animals and controls or between sexes during the dark cycle (Time: F(5.690, 79.65) = 2.900, *p=0.0146; Virus: F(1, 14) = 0.07592, p=0.7869; Virus_Total_: F(1, 14) = 0.2453, p=0.6281) or between sexes during the dark cycle (Sex: F(1, 14) = 3.448, #p=0.0845; Sex_Total_: F(1, 14) = 3.225, #p=0.0941). Within-sex comparisons reveal no effect of virus in males (Sidak’s multiple comparisons; **p_adj_=0.8862) or females (Sidak’s multiple comparisons; p_adj_=0.8994). Experiments conducted in the following light conditions: Yellow = light cycle only; Blue = dark cycle only; Red = during the light cycle but under red light conditions. #p<0.1, *p<0.05, **p<0.01, ***p<0.001, ****p<0.0001. Created in BioRender. Keefauver, T. (2026) <https://BioRender.com/z8o0u2s>
