## Supplementary figures and images for "Astrocyte molecular rhythm disruption in nucleus accumbens promotes increased binge-like drinking in mice"

### Figure S1

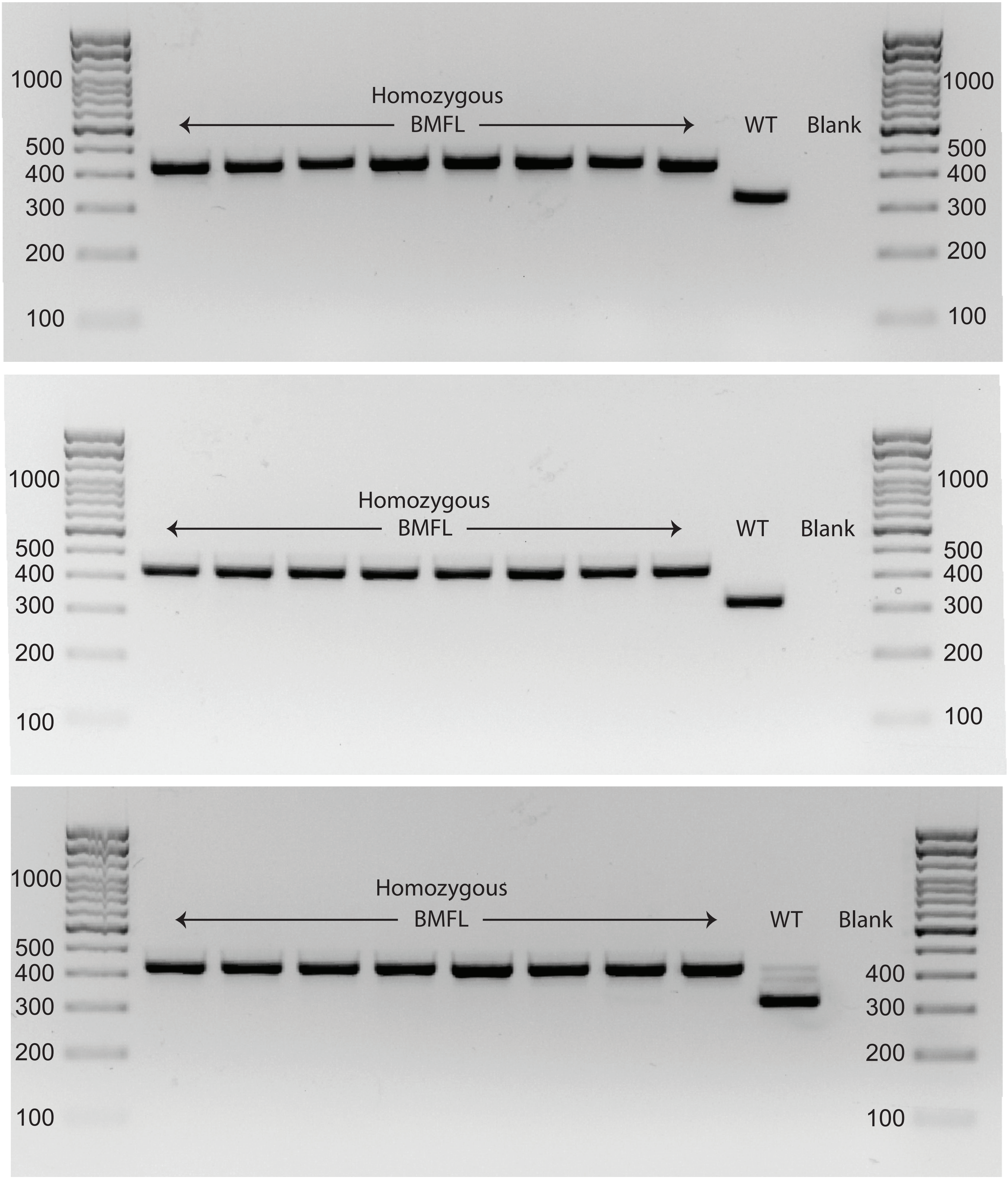

### Figure S2

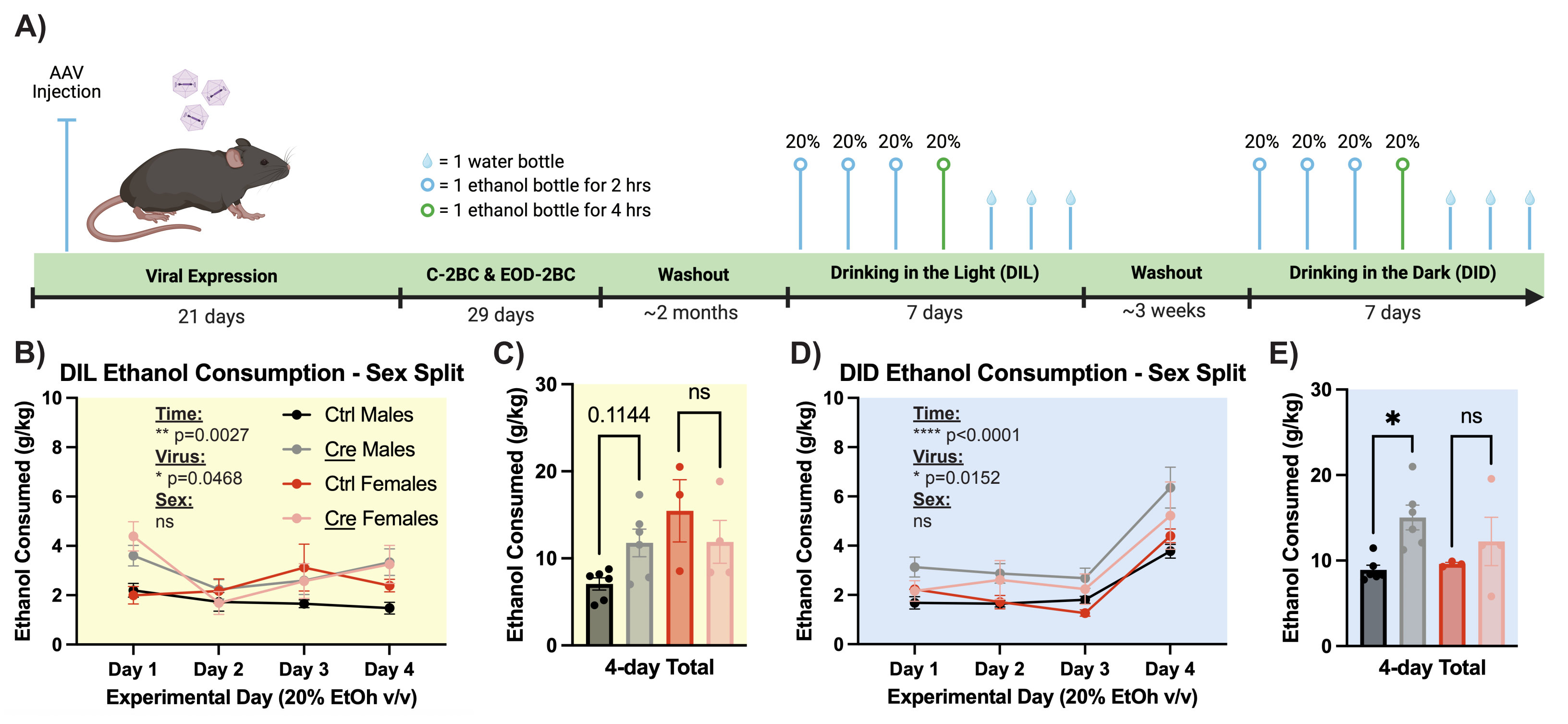

### Figure S3

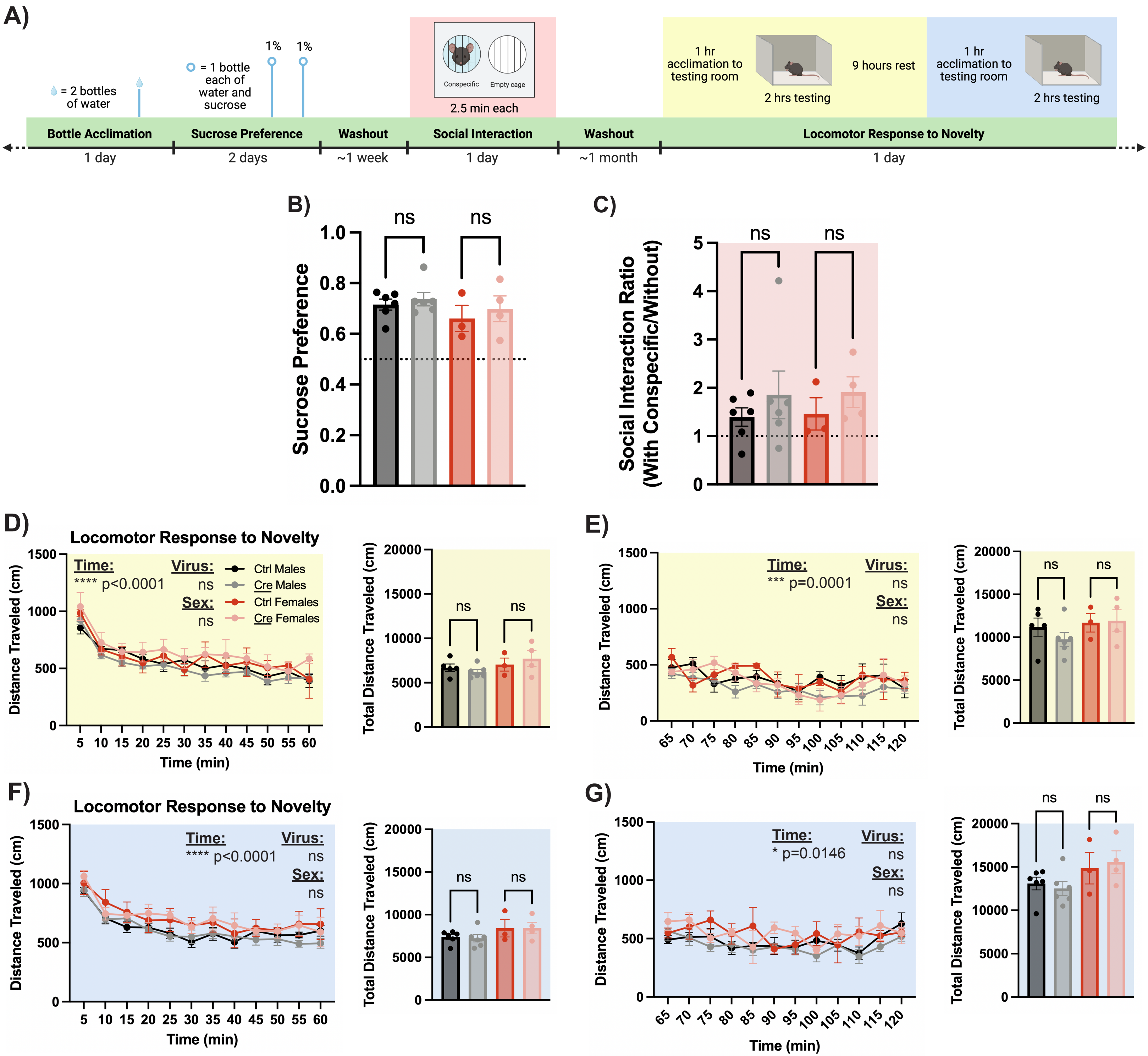
